# Immune cells promote paralytic disease in mice infected with enterovirus D68

**DOI:** 10.1101/2024.10.14.618341

**Authors:** Mikal A. Woods Acevedo, Jie Lan, Sarah Maya, Jennifer E. Jones, John V. Williams, Megan Culler Freeman, Terence S. Dermody

## Abstract

Enterovirus D68 (EV-D68) is associated with acute flaccid myelitis (AFM), a poliomyelitis-like illness causing paralysis in young children. However, mechanisms of paralysis are unclear, and antiviral therapies are lacking. To better understand EV-D68 disease, we inoculated newborn mice intracranially to assess viral tropism, virulence, and immune responses. Wild-type (WT) mice inoculated intracranially with a neurovirulent strain of EV-D68 showed infection of spinal cord neurons and developed paralysis. Spinal tissue from infected mice revealed increased levels of chemokines, inflammatory monocytes, macrophages, and T cells relative to controls, suggesting that immune cell infiltration influences pathogenesis. To define the contribution of cytokine-mediated immune cell recruitment to disease, we inoculated mice lacking CCR2, a receptor for several EV-D68-upregulated cytokines, or RAG1, which is required for lymphocyte maturation. WT, *Ccr2*^-/-^, and *Rag1*^-/-^ mice had comparable viral titers in spinal tissue. However, *Ccr2*^-/-^ and *Rag1*^-/-^ mice had significantly less paralysis relative to WT mice. Consistent with impaired T cell recruitment to sites of infection in *Ccr2*^-/-^ and *Rag1*^-/-^ mice, antibody-mediated depletion of CD4^+^ or CD8^+^ T cells from WT mice diminished paralysis. These results indicate that immune cell recruitment to the spinal cord promotes EV-D68-associated paralysis and illuminate new targets for therapeutic intervention.

**Graphical Abstract:** 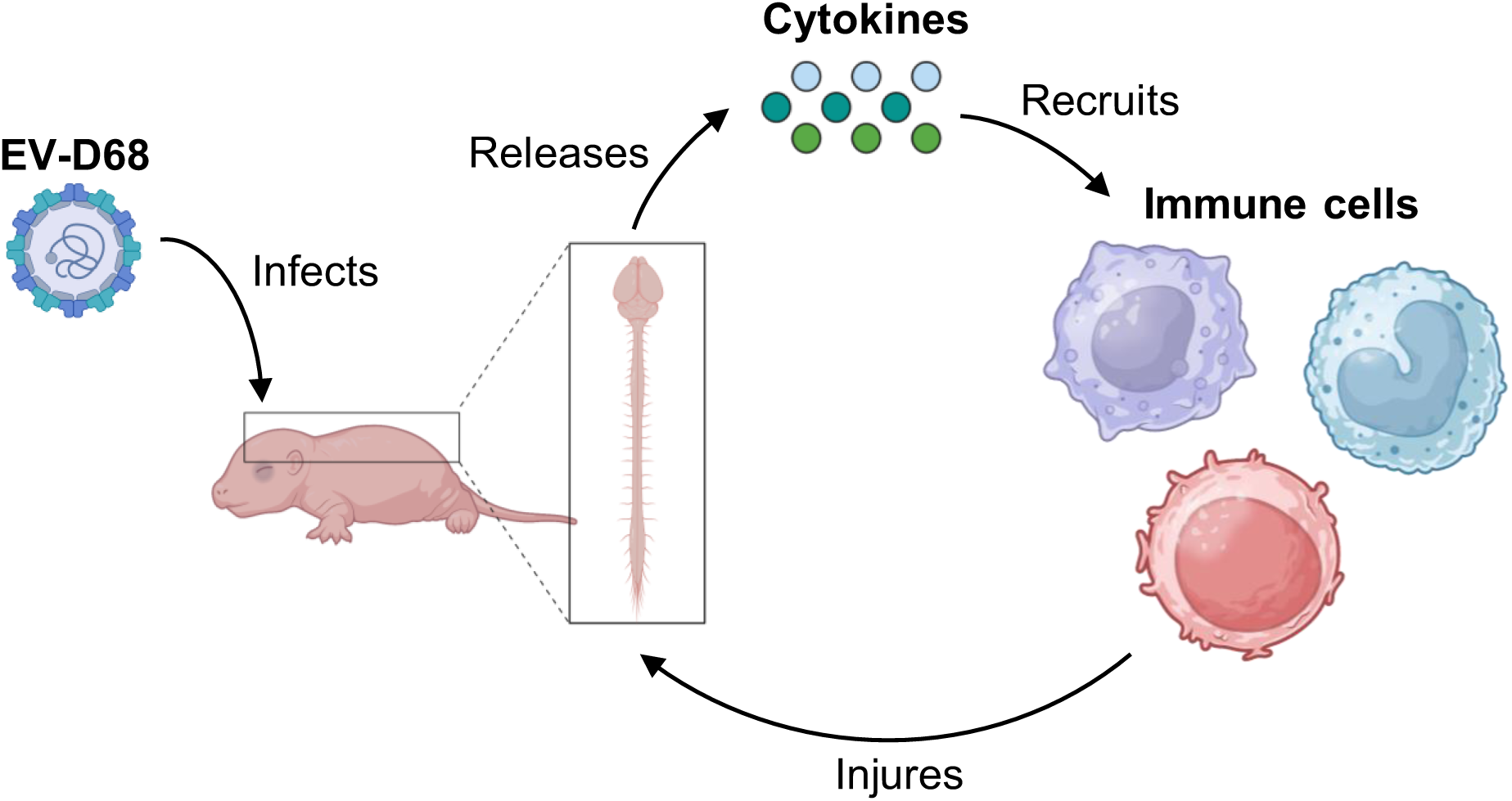

## Introduction

Enteroviruses cause a wide spectrum of disease in humans, including acute flaccid myelitis (AFM), a poliomyelitis-like paralytic condition that occurs primarily in children (1,2). AFM is thought to result from injury to spinal cord motor neurons (3,4), although the mechanisms of cell killing are unclear. EV-D68 was first detected in 1962 in children with pneumonia (5) and is considered a reemergent pathogen after its association with AFM. The CDC began tracking AFM outbreaks in 2014 (6,7), but EV-D68 has been associated with paralysis as early as 2008 (3,8). There are currently no targeted treatments for EV-D68 infections or AFM (9). Therefore, there is an urgent need to define mechanisms by which EV-D68 causes disease.

Other neurotropic viruses, such as poliovirus, cause limb paralysis by inducing apoptosis in spinal cord motor neurons (10). In human spinal cord organoids (hSCOs), which lack immune cells, EV-D68 replicates but causes minimal cell death relative to other enteroviruses (11), suggesting that EV-D68 replication is not the sole mediator of neuronal cell death. Cerebral spinal fluid (CSF) obtained from persons with AFM rarely contains evidence of a pathogen (12). However, CSF from those with AFM is often enriched for enterovirus-specific antibodies (13,14). Post-mortem studies of a child who developed flaccid paralysis yielded EV-D68 RNA in the CSF and identified EV-D68 capsid protein and RNA, CD68^+^ macrophages, and CD8^+^ T cells in the spinal cord (3,8). Furthermore, spinal cord sections from this child stained negative for caspase-3, but positive for perforin, suggesting that the child’s immune response, and not virus-induced apoptosis, contributed to the disease (3). Immune cells can exacerbate certain virus-induced diseases. For example, CD8^+^ T cells contribute to Zika virus-associated paralysis (15) and LCMV-related mortality in mice (16). Mature lymphocytes also are implicated in damaging myelin after spinal cord injury, which restricts recovery (17). Uncovering mechanisms by which EV-D68 and the subsequent immune response contributes to disease progression could potentially lead to the identification of targets for therapeutic intervention.

Several *in vitro* and *ex vivo* models are available to study EV-D68 neural infection (18). Contemporary neurotropic EV-D68 strains, but not historical non-neurotropic strains, efficiently replicate in human neuronal cell lines (19). Viral replication efficiency in human neuroblastoma cells is attributable to sequence polymorphisms in the viral capsid (20). In murine organotypic brain slice cultures, EV-D68 infects Nissl-stained neurons (21). In primary rat cortical neurons, EV-D68 infects both excitatory glutamatergic and inhibitory GABAergic neurons (22). Cultivated human B cells and dendritic cells, but not CD4^+^ or CD8^+^ T cells, stain positive for EV-D68 capsid protein (23), suggesting that both neurons and certain immune cell subsets can be infected by EV-D68 in humans. However, while *in vitro* and *ex vivo* models serve as tractable systems to study how EV-D68 infects cells, they do not recapitulate how EV-D68 disease develops in a complex host environment.

Animal models of EV-D68 disease reproduce important aspects of infection and disease (18). In newborn mice, EV-D68 infection causes an AFM-like illness (9). As mice age, they become resistant to EV-D68-associated AFM (24). Additionally, the route of inoculation and viral dose dictates the efficiency by which EV-D68 causes paralysis, with intracranial inoculations being one of the most neuropathogenic routes (9,18). Thus, neonatal mice serve as useful experimental models to study mechanisms by which EV-D68 causes AFM.

Here, we examined the effect of host immunity on EV-D68 replication in the central nervous system (CNS) and how it influences disease development in newborn mice. We observed an EV-D68 strain-specific cytokine response in both hSCOs and in newborn mice, which had robust immune cell recruitment to the spinal cord. Additionally, we observed that mice lacking functional immune cell recruitment or mature lymphocytes had diminished disease relative to immunocompetent mice. Antibody-mediated depletion of CD8^+^ or CD4^+^ T cells resulted in significant protection against EV-D68-associated AFM relative to isotype control antibody. Collectively, these findings suggest that immune cells recruited to the spinal cord promote development of EV-D68-associated AFM.

## Results

### EV-D68 strains differ in virulence and immune responses in immunocompetent neonatal wildtype mice

To understand how EV-D68 causes paralysis, we recovered the prototype USA/Fermon strain, non-virulent strain US/MO/14-18949 (MO49) (25), and virulent strain US/IL/14-18952 (IL52) (25) from cDNA clones. We inoculated 3-day-old C57BL6/J wildtype (WT) mice intracranially (i.c.) with 10^5^ PFU of each strain. At 3 days post-inoculation (dpi), we resected brains and spinal columns from infected mice and determined viral loads in homogenized tissues by plaque assay (Figure 1A). Only mice inoculated with EV-D68 IL52 had appreciable viral titers in spinal tissue, which prompted us to examine viral replication kinetics of this strain in brain and spinal tissue by determining titers at 1, 3, and 5 dpi (Figure 1B). Viral loads at the site of inoculation were detectable at 1 dpi, whereas replication in the spinal column was sustained. To define sites of EV-D68 infection in the spinal column, we stained spinal cord sections for the EV-D68 VP1 capsid protein and observed viral antigen associated with NeuN-marked neurons in infected animals (Figure 1C). Additionally, EV-D68 IL52 was the only strain tested that caused paralysis in WT mice (Figure 1D), as has been reported previously (9,25). In contrast to AFM in humans (26), there was no observable pattern in the distribution of limbs affected by paralysis in infected mice (Supplemental Figure 1A). Most EV-D68 IL52-inoculated mice that did not develop paralysis had detectable EV-D68-neutralizing antibody titers by day 14 post-inoculation (Figure 1E), indicating that paralysis following infection was not uniform. Collectively, these results demonstrate that some strains of EV-D68 replicate, cause disease, and elicit neutralizing antibodies in neonatal WT mice inoculated intracranially.

**Figure 1.**
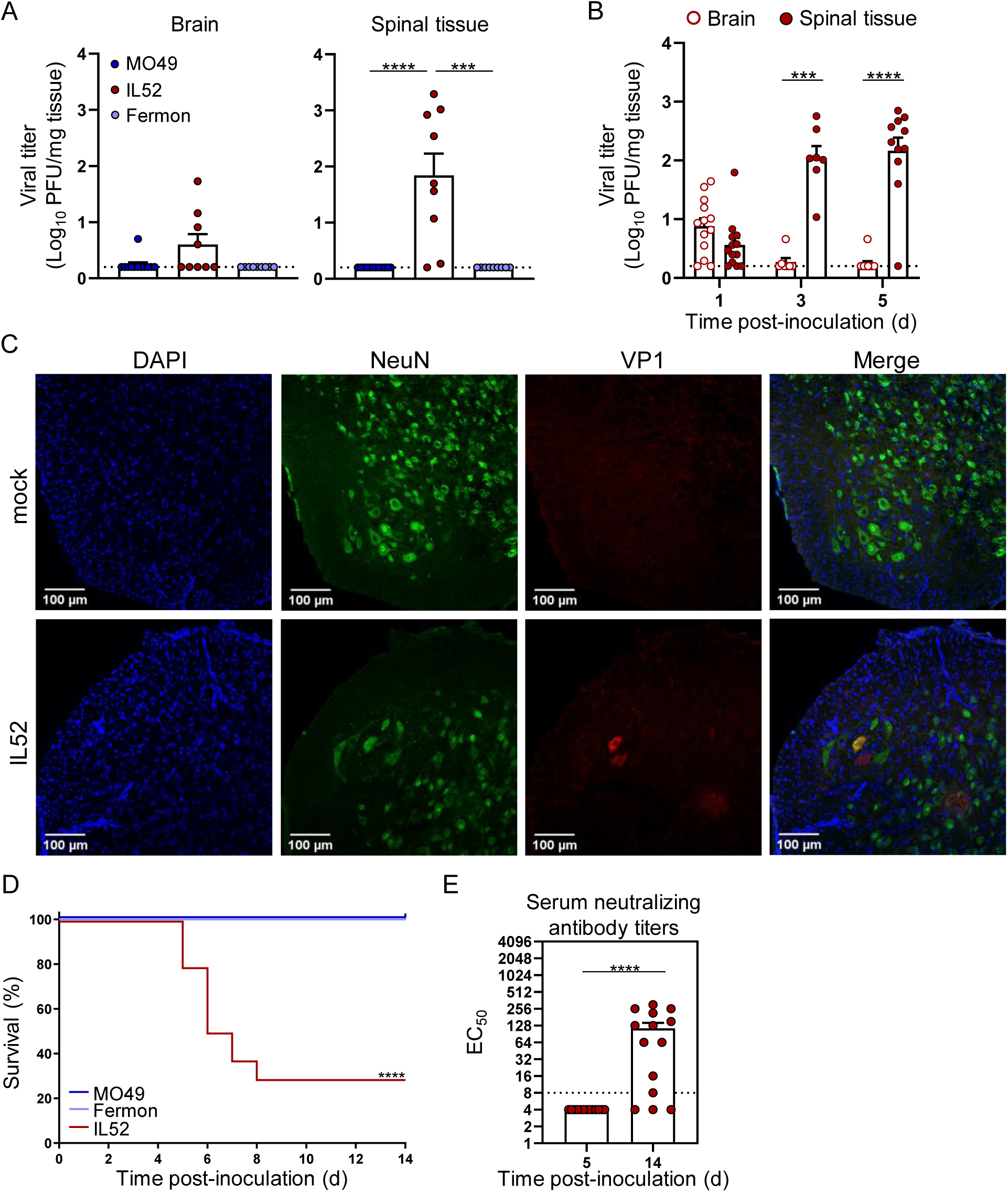
EV-D68 US/IL/14-18952 infects and causes paralysis in immunocompetent neonatal mice. Three-day-old WT mice were inoculated i.c. with PBS (mock) or 10^5^ PFU of EV-D68 USA/Fermon, US/MO/14-18949 (MO49), or US/IL/14-18952 (IL52). (**A**) Brain and spinal tissue were resected at 3 dpi from EV-D68-inoculated mice, and viral titers were determined by plaque assay. (**B**) Brain and spinal tissue were resected at 1, 3, or 5 dpi from IL52-inoculated mice, and viral titers were determined by plaque assay. (**C**) Spinal columns were resected at 5 dpi from IL52-inoculated paralyzed mice or day-matched mock-inoculated controls. Samples were processed for immunohistochemistry staining. DAPI, blue; NeuN, green; EV-D68 VP1, red. Scale bar, 100 μM. (**D**) Inoculated mice were monitored daily and euthanized upon signs of paralysis. N = 12-27 mice per group. (**E**) Neutralizing antibody titers in sera collected at 5 or 14 dpi from IL52-inoculated mice. Dotted lines indicate the limit of detection. Data are representative of 2-3 independent experiments. Each symbol represents an individual mouse. Error bars indicate mean ± SEM. Mann-Whitney test (**A**, **B**, and **E**) or log-rank test (**D**): *, *P* ≤ 0.05; **, *P* ≤ 0.01; ***, *P* ≤ 0.001; ****, *P* ≤ 0.0001.

### EV-D68 strains differ in cytokine responses in spinal tissue of WT mice

To characterize the host response to EV-D68, we conducted multianalyte Luminex-based profiling of 31 pro-inflammatory cytokines in spinal tissue of WT mice that were inoculated with PBS as a control or EV-D68 strains MO49 or IL52 (Figure 2A). At 3 dpi, a subset of cytokines were elevated in spinal tissue from mice inoculated with either virus strain. However, mice inoculated with the neurovirulent strain IL52 had significantly higher levels of CCL7 and CCL12, and to a lesser extent CCL2, relative to mice inoculated with EV-D68 MO49 (Figure 2B). These results indicate that a neurovirulent EV-D68 strain elicits a robust cytokine response in spinal tissue of WT mice.

**Figure 2.**
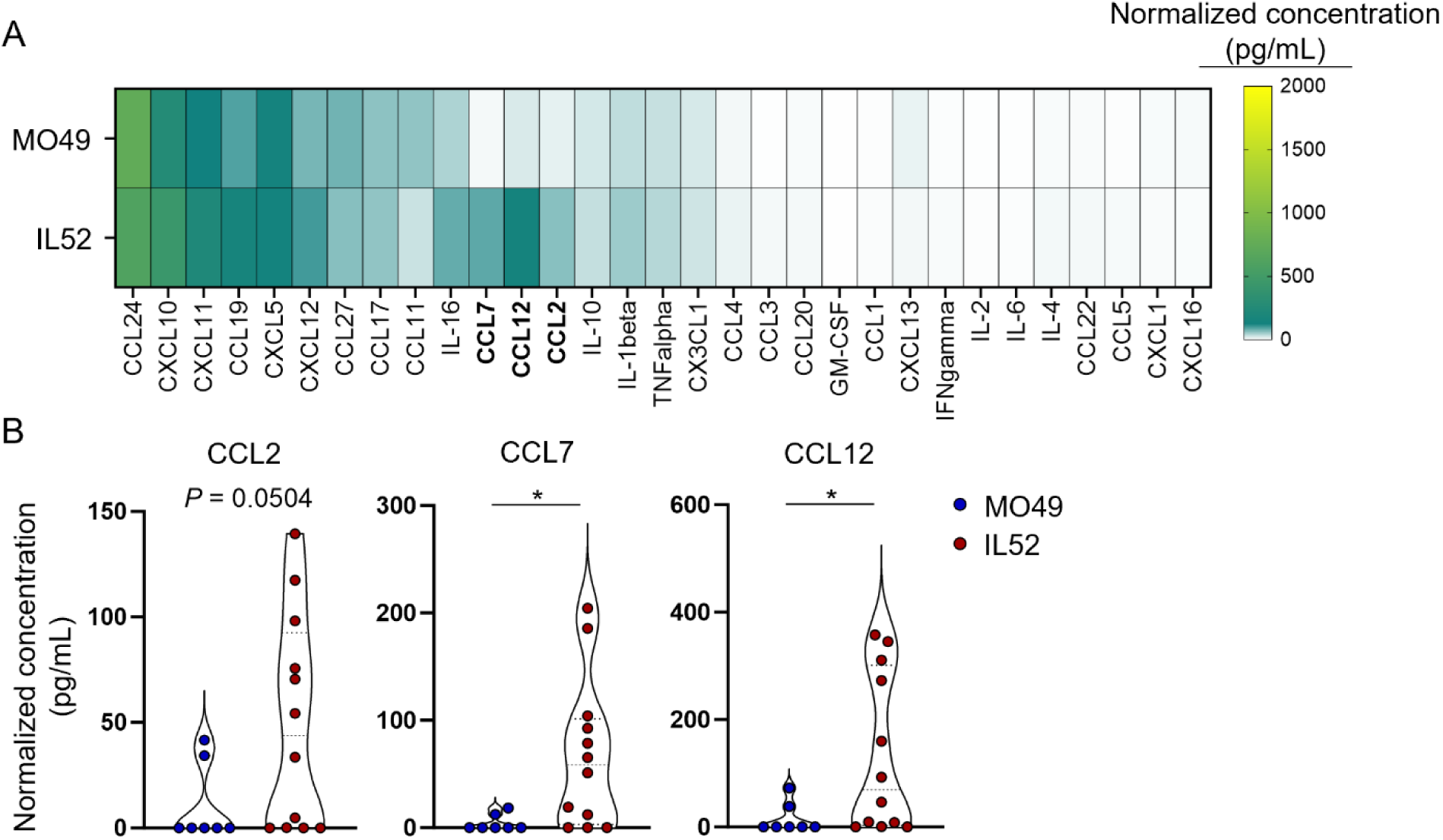
Cytokines in spinal tissue of EV-D68-inoculated mice. Three-day-old WT mice were inoculated i.c. with MO49, IL52, or PBS (mock). Spinal tissue was resected 3 dpi and analyzed by Luminex protein assay. (**A**) Cytokine concentrations are presented as mean values in pg/mL from 7-12 mice and shown as a heatmap normalized to mock-inoculated mice. (**B**) CCL2, CCL7, and CCL12 concentrations normalized to the corresponding concentrations in mock-inoculated mice are shown. Data are representative of 2-3 independent experiments. Each symbol represents an individual mouse. Mann-Whitney test (**B**): *, *P* ≤ 0.05.

### Neurovirulent EV-D68 infects human spinal cord organoids and elicits a cytokine response that mimics the cytokine response in mice

EV-D68 strains that differ in virulence differ in cytokine responses in neonatal mice, but it is unclear what cells respond to infection and dominate the response. To determine how a multicellular organoid responds to viral infection in the absence of immune cells and to assess whether the cytokine response in this system mimics that observed in inoculated mice, we used a human-derived 3-dimensional spinal cord organoid (3-DiSC hSCO) system to characterize the cytokine response to EV-D68 infection. We inoculated pools of 8-12 organoids with PBS (mock) or EV-D68 strains MO49 or IL52 and monitored infection and cytokine levels. Consistent with the trends in viral load in neural tissue of mice, we observed robust VP1 staining in organoids inoculated with IL52 but not MO49 at 3 dpi (Figure 3A). We next examined virus-mediated cytokine profiles in 3DiSC hSCO supernatants harvested at 3 dpi (Figure 3B). Inoculation with EV-D68 IL52 resulted in cytokine induction similar to that in mice. Together, these results indicate that neurovirulent EV-D68 IL52 infects 3-DiSC hSCO and induces a cytokine profile comparable to that in murine spinal tissue.

**Figure 3.**
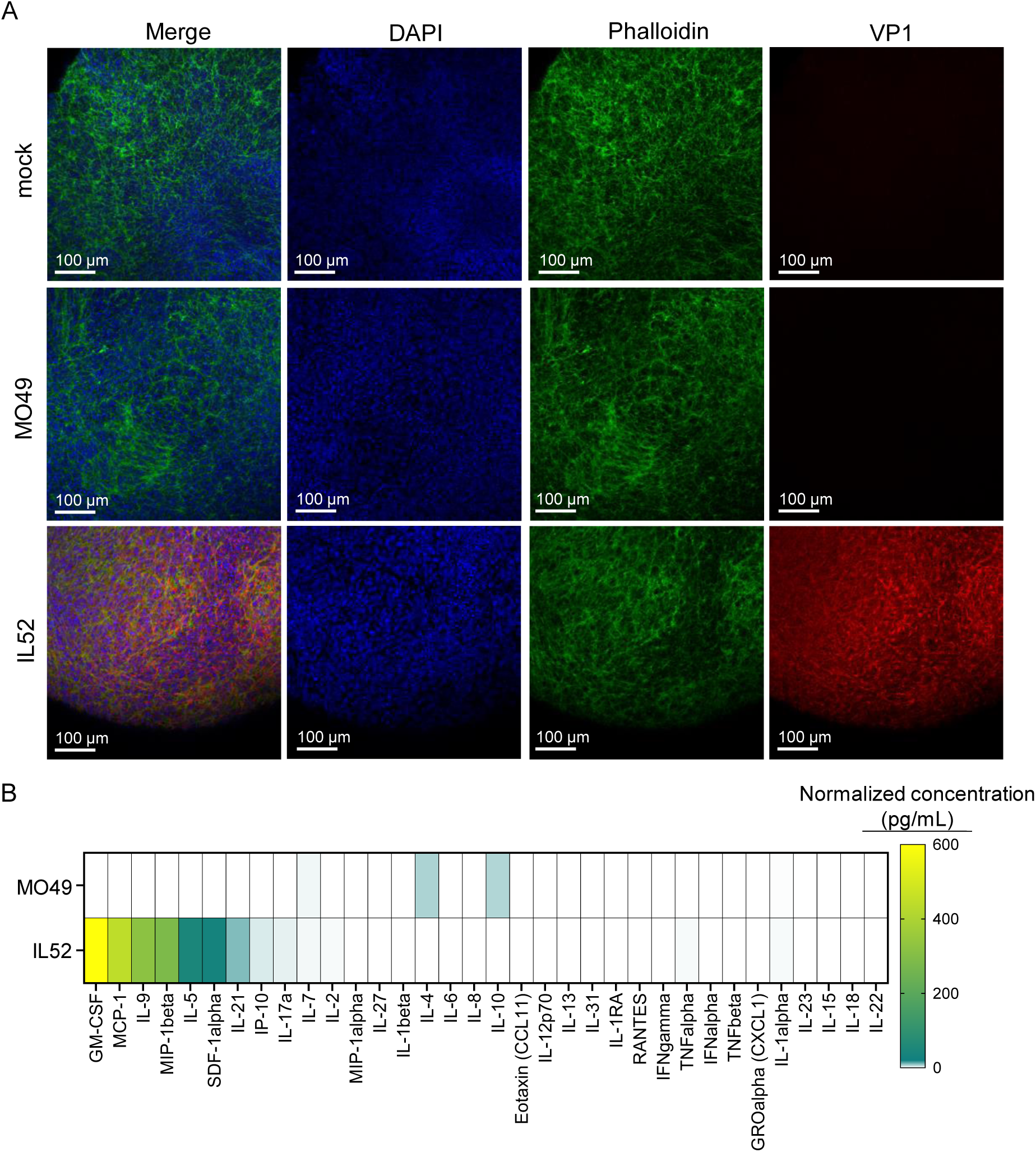
Neurovirulent EV-D68 efficiently infects human spinal cord organoids. Pools of 8-12 hSCOs at 14 days post-differentiation were either mock-inoculated (PBS) or inoculated with 10^5^ PFU of EV-D68 MO49 or IL52. (**A**) At 3 dpi, whole organoids were processed for immunofluorescence. DAPI, blue; phalloidin, green; EV-D68 VP1, red. (**B**) hSCO supernatants were collected at 3 dpi and analyzed by Luminex protein assay. Protein levels are presented as mean values in pg/mL from 2-3 independent samples and shown as a heatmap normalized to mock-infected organoids.

### Spinal cords of paralyzed mice have increased immune cell populations

Since spinal tissue of mice inoculated with EV-D68 IL52 produced increased levels of chemoattractant cytokines relative to those inoculated with PBS or EV-D68 MO49, we hypothesized that neurovirulent EV-D68 infection leads to immune cell recruitment to the spinal cord. Furthermore, CD8^+^ T cells and CD68^+^ macrophages were observed in spinal cord sections demonstrating enterovirus antigen from a patient who died of AFM (8), suggesting that cellular immunity contributes to AFM pathogenesis. To assess the immune cell environment in the spinal cord of paralyzed mice, we examined spinal cord single-cell suspensions using a multiplexed 27-cell-type flow cytometry panel (Supplemental Table 1 and Supplemental Figure 2). Spinal cords were resected from mice inoculated with EV-D68 IL52 that displayed signs of paralysis or day-matched controls inoculated with PBS or EV-D68 MO49 and analyzed by flow cytometry (Figure 4A). There were significant increases in the numbers of total T cells, CD4^+^ T cells, CD8^+^ T cells, total macrophages, M1 macrophages, total monocytes, and inflammatory monocytes in the spinal cord of EV-D68 IL52-inoculated mice relative to those inoculated with PBS or EV-D68 MO49. There also was an increased percentage of CD4^+^ T cells, CD8^+^ T cells, total macrophages, M1 macrophages, and inflammatory monocytes in the spinal cords of EV-D68 IL52-inoculated mice relative to controls (Figure 4B). Other immune cell populations varied to a more modest extent or did not differ significantly between the groups (Supplemental Figure 3A-B). These results suggest that infection with a neurovirulent strain of EV-D68 leads to an increase in a variety of immune cell types in the spinal cord.

**Figure 4.**
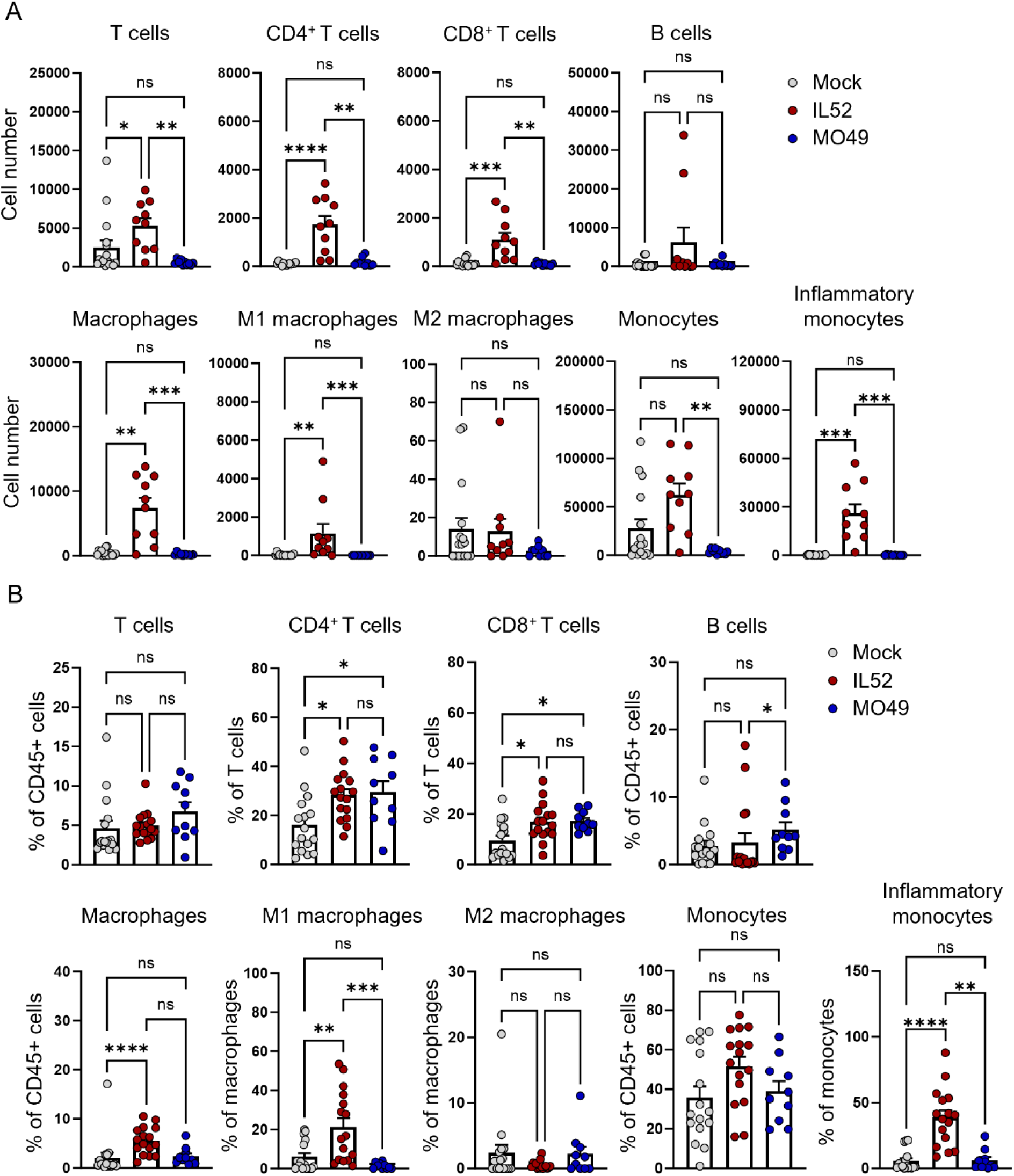
Mice inoculated with neurovirulent EV-D68 have altered populations of immune cells in the spinal cord. Three-day-old WT mice were inoculated i.c. with PBS (mock) or EV-D68 MO49 or IL52. Spinal cords were resected from paralyzed mice inoculated with IL52 or day-matched mice inoculated with MO49 or PBS. Single-cell suspensions were prepared, stained, and analyzed by flow cytometry. (**A**) Numbers and (**B**) percentages of selected cell types are shown. Data are representative of 2-4 independent experiments. Each symbol represents an individual mouse. Error bars indicate mean ± SEM. Mann-Whitney

### CCR2-dependent immune cell recruitment influences paralysis frequency

As EV-D68 IL52-inoculated mice had higher levels of chemoattractant cytokines and increased immune cell infiltrates relative to controls, we hypothesized that cytokine-induced immune cell recruitment influences EV-D68-mediated paralysis. To determine whether lymphocytes recruited to the spinal cord regulate EV-D68 disease, we assessed EV-D68 replication and virulence in mice lacking C-C chemokine receptor type 2 (*Ccr2^-/-^*), which mediates immune cell recruitment and, importantly, is a common receptor used by CCL2, CCL7, and CCL12 (27), which are upregulated by neurovirulent EV-D68 IL52. We inoculated WT or *Ccr2^-/-^* mice with EV-D68 IL52, resected brain and spinal tissues at 1, 3, and 5 dpi, and quantified viral loads by plaque assay. There were no statistically significant differences in viral loads in spinal tissue of WT and *Ccr2^-/-^* mice and only a modest increase in viral loads in brain tissue of *Ccr2^-/-^* mice at 3 dpi (Figure 5A). Surprisingly, despite similar viral loads, *Ccr2^-/-^* mice developed less paralysis relative to WT mice (Figure 5B) and did not manifest an observable pattern in paralyzed limb distribution (Supplemental Figure 1B). To define the immune response to EV-D68 IL52 infection in WT and *Ccr2^-/-^* mice, we conducted multianalyte Luminex-based assays for 31 pro-inflammatory cytokines in spinal tissue of infected mice (Figure 5C). There were increased levels of CCL2 but not CCL7 or CCL12 in the spinal cords of *Ccr2^-/-^* mice relative to WT mice (Figure 5D). Collectively, these data suggest that CCR2-deficient mice are significantly protected against EV-D68 disease.

**Figure 5.**
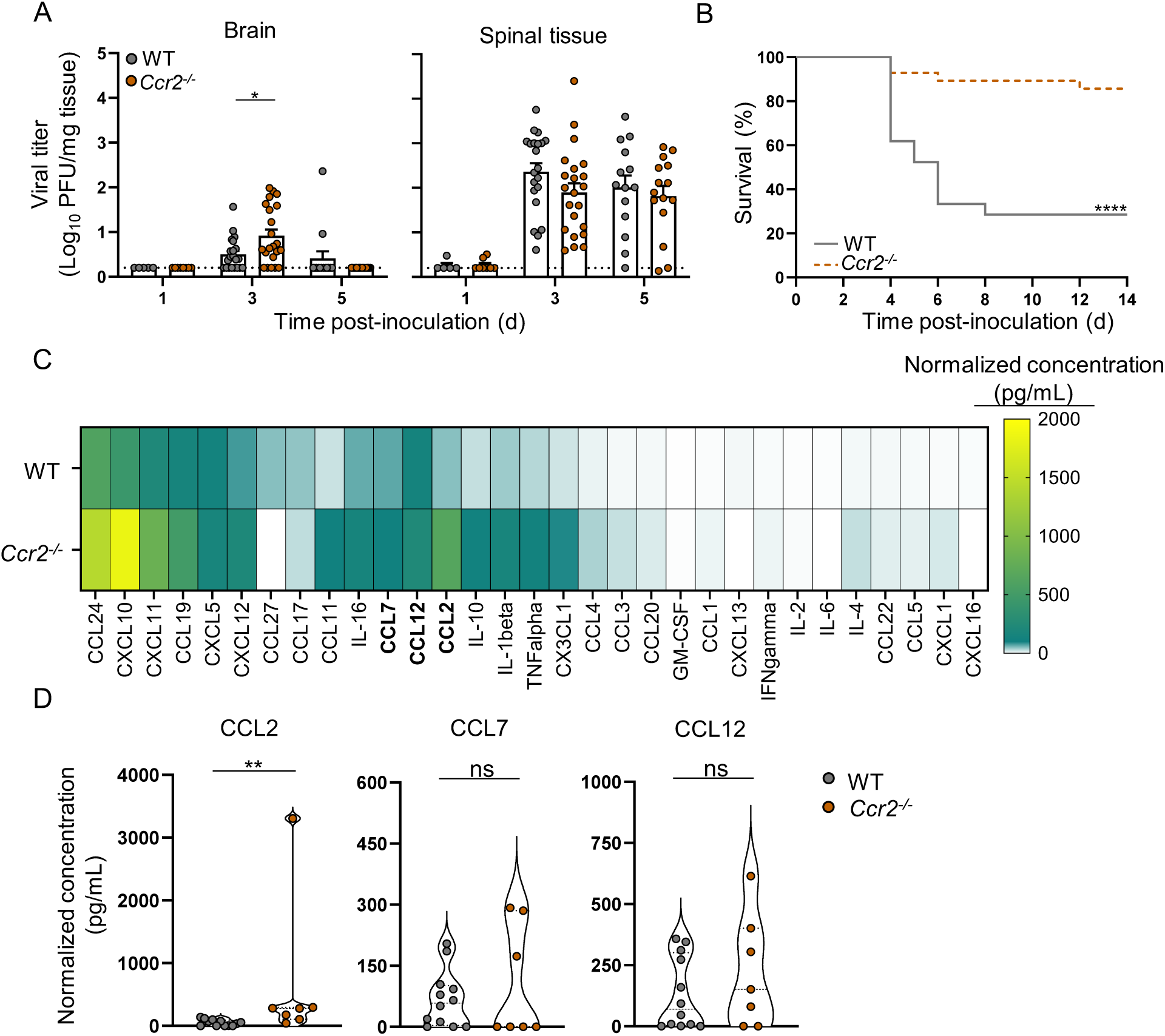
Mice lacking CCR2 develop less paralysis relative to WT mice following inoculation with neurovirulent EV-D68. Three-day-old WT or *Ccr2^-/-^* mice were inoculated i.c. with PBS (mock) or EV-D68 IL52. (**A**) Brain and spinal tissue of virus-inoculated mice were resected at 1, 3, and 5 dpi, and viral titers were determined by plaque assay. Dotted lines indicate the limit of detection. (**B**) Virus-inoculated mice were monitored daily and euthanized upon signs of paralysis. N = 21-28 mice per group. (**C**) Spinal tissue was resected 3 dpi and analyzed by Luminex protein assay. Data from WT mice are the same as those from IL52-inoculated mice in Figure 1F, as these experiments were conducted concurrently. Cytokine concentrations are presented as mean values in pg/mL from 3-12 mice and shown as a heatmap normalized to mock-inoculated mice. (**D**) Levels of CCL2, CCL7, and CCL12 are shown. Data are representative of 2-3 independent experiments. Each symbol represents an individual mouse. Error bars indicate mean ± SEM. Mann-Whitney test (**A and D**) or log-rank test (**B**): *, *P* ≤ 0.05; **, *P* ≤ 0.01; ****, *P* ≤ 0.0001.

### Immune cell recruitment following EV-D68 infection is impaired in Ccr2^-/-^ mice

Since *Ccr2^-/-^* mice develop significantly less paralysis following EV-D68 infection than WT mice, we sought to define differences in the spinal cord immune cell profile of *Ccr2^-/-^* and WT mice using flow cytometry. Spinal cords were resected from EV-D68 IL52-inoculated WT mice that displayed signs of paralysis or day-matched infected *Ccr2^-/-^* mice. Spinal cords of WT mice had increased numbers of total T cells, CD4^+^ T cells, CD8^+^ T cells, total macrophages, and M1 macrophages relative to *Ccr2^-/-^* mice (Figure 6A). Additionally, there was an increased percentage of total macrophages in the spinal cords of WT mice relative to *Ccr2^-/-^* mice (Figure 6B). Other immune cell populations varied to a more modest extent or did not differ significantly between WT and *Ccr2^-/-^* mice (Supplemental Figure 4A-B). We conclude that CCR2-deficient mice have impaired immune cell recruitment to the spinal cord relative to WT mice following EV-D68 infection.

**Figure 6.**
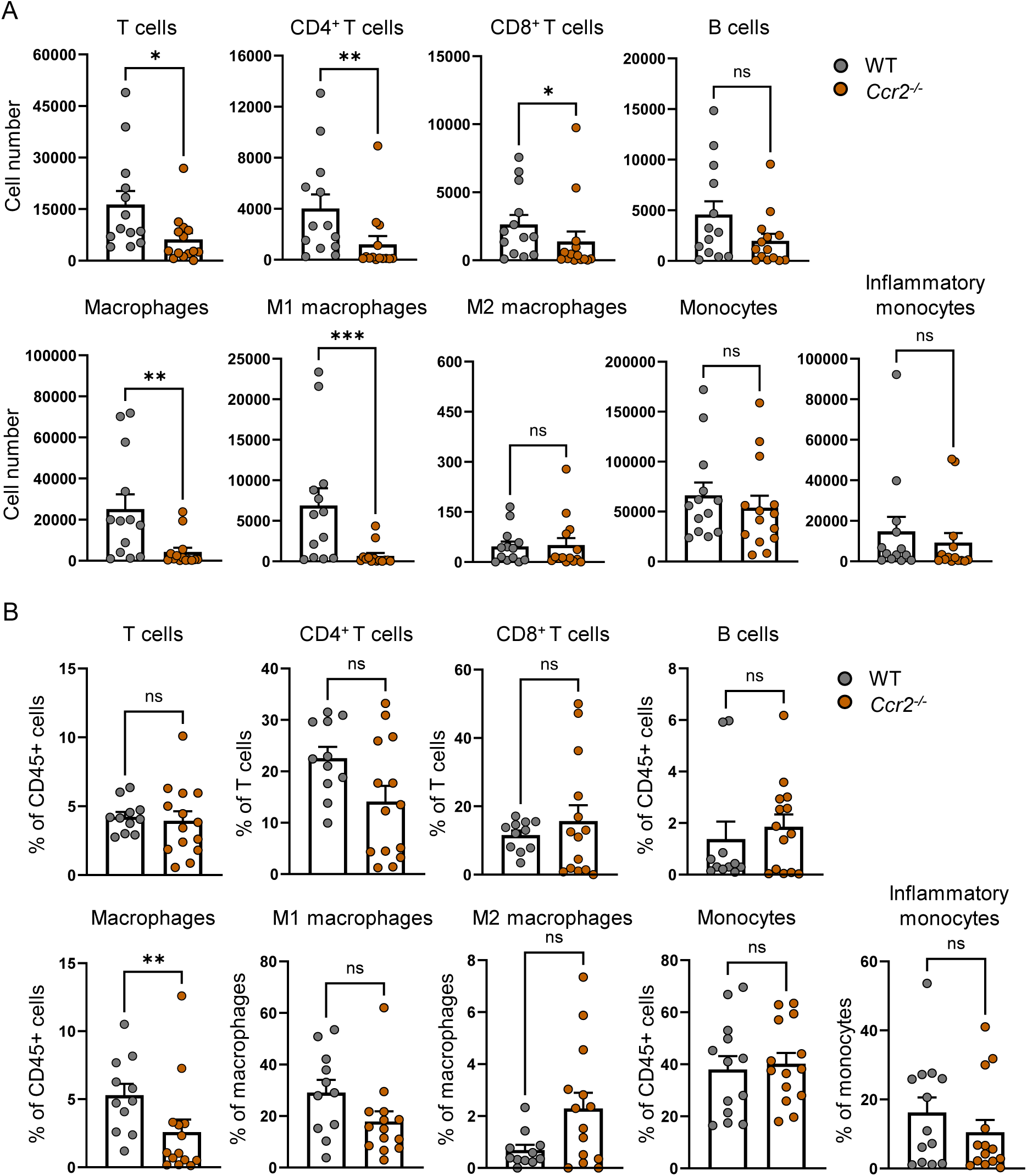
Loss of CCR2 results in altered immune cell recruitment following neurovirulent EV-D68 inoculation. Three-day-old WT or *Ccr2^-/-^* mice were inoculated i.c. with EV-D68 IL52. Spinal cords were resected at 5-6 dpi from paralyzed WT mice or day-matched paralyzed and non-paralyzed *Ccr2^-/-^* mice. Single-cell suspensions were prepared, stained, and analyzed by flow cytometry. (**A**) Numbers and (**B**) percentages of selected cell types are shown. Each symbol represents an individual mouse. Error bars indicate mean ± SEM. Mann-Whitney test: *, *P* ≤ 0.05; **, *P* ≤ 0.01 ***, *P* ≤ 0.001; ns = not significant.

### Paralysis frequency following EV-D68 infection is diminished in Rag1-deficient mice

In our experiments, T cells are robustly recruited to the spinal cord of WT mice following infection with EV-D68. Since T cells contribute to disease severity following infection with other neurovirulent viruses (15,16), we hypothesized that immune cells contribute to acute EV-D68 disease in mice. To determine whether B and T lymphocytes affect EV-D68 replication and virulence, we inoculated either WT mice or *Rag1^-/-^* mice, which lack mature B and T lymphocytes, with EV-D68 IL52 and quantified viral loads at 3 dpi by plaque assay (Figure 7A). There was a modest decrease in viral load in the brains of *Rag1^-/-^* mice relative to WT mice, but there was no significant difference in viral load in spinal tissue. Strikingly, *Rag1^-/-^* mice inoculated with EV-D68 IL52 developed significantly less paralysis compared with WT mice (Figure 7B), and no infected *Rag1^-/-^* mice had multi-limb paralysis (Supplemental Figure 1B). *Rag1^-/-^* mice had no detectable neutralizing antibodies at 14 dpi, consistent with the absence of lymphocytes in these animals (Supplemental Figure 5). However, *Rag1^-/-^* mice had detectable virus in spinal tissue at 14 dpi, whereas WT and *Ccr2^-/-^* mice had no detectable virus at this timepoint (Supplemental Figure 5). These results indicate that while lymphocytes aid in clearing virus they also promote EV-D68 disease.

**Figure 7.**
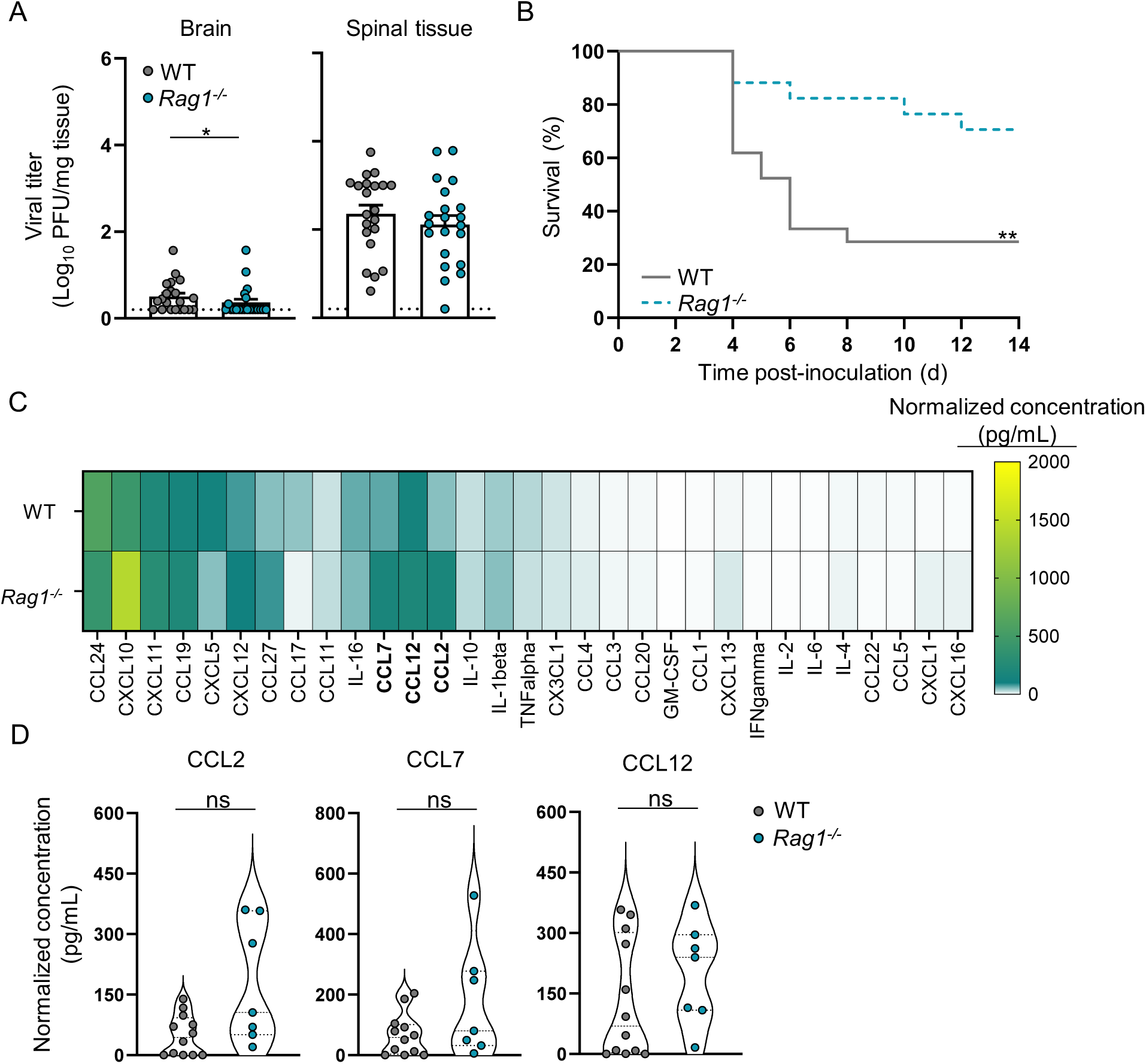
Mice lacking mature B and T cells have diminished EV-D68-induced paralysis relative to WT mice. Three-day-old WT or *Rag1^-/-^* mice were inoculated i.c. with PBS (mock) or EV-D68 IL52. (**A**) Brain and spinal tissue were resected at 3 dpi, and viral titers were determined by plaque assay. Data from WT mice are the same as those presented in Figure 5A, as these experiments were conducted concurrently. Dotted lines indicate the limit of detection. (**B**) Virus-inoculated mice were monitored daily and euthanized upon signs of paralysis. N = 16-21 mice per group. Data from WT mice are the same as those presented in Figure 5C, as these experiments were conducted concurrently. (**C**) Spinal tissue was resected 3 dpi and analyzed by Luminex protein assay. Data from WT mice are the same as those presented in Figure 2A, as these experiments were conducted concurrently. Cytokine concentrations are presented as mean values in pg/ml from 7-12 mice and shown as a heatmap normalized to mock-inoculated mice. (**D**) Levels of CCL2, CCL7, and CCL12 are shown. Data are representative of 2-3 independent experiments. Each symbol represents an individual mouse. Error bars indicate mean ± SEM. Mann-Whitney test (**A and D**) or log-rank test (**B**): *, *P* ≤ 0.05; **, *P* ≤ 0.01; ns = not significant.

To characterize the cytokine response elicited by EV-D68 in WT and *Rag1^-/-^* mice, we conducted multianalyte Luminex-based assays for 31 pro-inflammatory cytokines in spinal tissue of mice inoculated with EV-D68 IL52 (Figure 7C). There were no statistically significant differences in the levels of CCL2, CCL7, or CCL12 in the spinal tissue of *Rag1^-/-^* and WT mice following inoculation with EV-D68 IL52 (Figure 7D). These observations suggest that lymphocytes are not required for the cytokine response to EV-D68 in the spinal cord.

To define the types of immune cells recruited into spinal tissue of *Rag1^-/-^* mice inoculated with EV-D68, we used flow cytometry to monitor 27 immune cell subtypes. The spinal tissue of *Rag1^-/-^* mice inoculated with EV-D68 IL52 had increased numbers of total macrophages, M1 macrophages, and inflammatory monocytes relative to spinal tissue of *Rag1^-/-^* mice inoculated with PBS as a control (Supplemental Figure 6A). Additionally, there was an increased percentage of CD4^+^ T cells, total macrophages, M2 macrophages, total monocytes, and inflammatory monocytes in spinal tissue of EV-D68 IL52-inoculated *Rag1^-/-^* mice relative to those inoculated with PBS (Supplemental Figure 6B). Other immune cell types varied to a more modest extent or did not differ significantly between *Rag1^-/-^* mice inoculated with EV-D68 or PBS (Supplemental Figure 7). These data suggest that mice lacking mature B and T lymphocytes have an altered immune cell population following EV-D68 IL52-inoculation.

### Antibody-mediated T cell depletion attenuates EV-D68-mediated paralysis

Since B cells are not efficiently recruited to the spinal cord following EV-D68 infection, we anticipated that the enhanced survival of *Rag1^-/-^* mice was attributable to the absence of T cells. To determine whether T cells influence EV-D68 disease, we administered antibodies specific for CD4^+^ or CD8^+^ to WT mice prior to inoculation with EV-D68 IL52 and assessed differences in disease penetrance (Figure 8A). Administration of antibodies against CD8^+^ T cells, and to a lesser extent CD4^+^ T cells, significantly protected mice from paralysis relative to those pretreated with an isotype antibody control (Figure 8B). These results indicate that T cells, specifically CD8^+^ T cells, promote EV-D68 paralysis and potentially identify new therapeutic targets against EV-D68-associated AFM.

**Figure 8.**
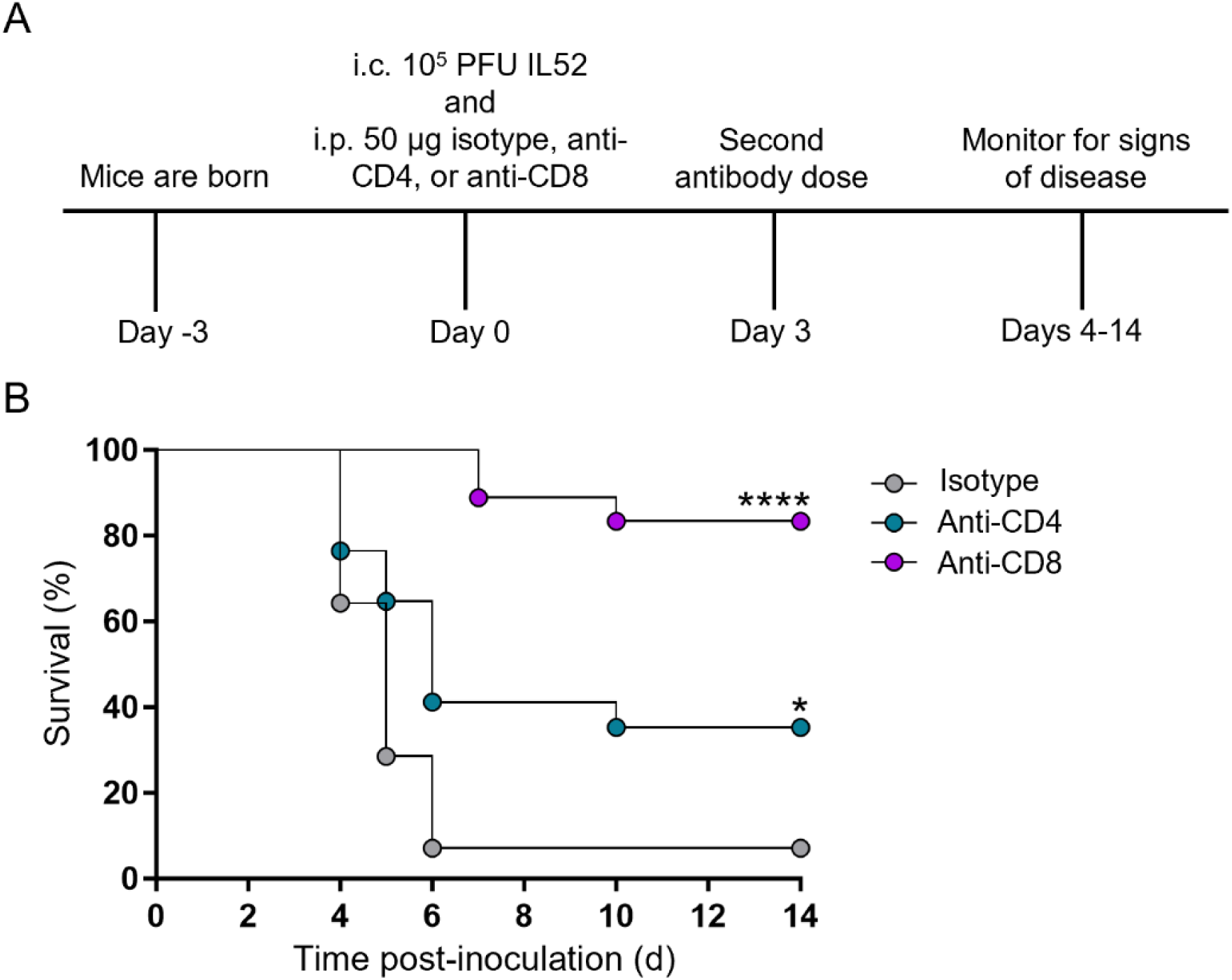
Antibody-mediated depletion of T cells protects mice from EV-D68-mediated paralysis. Three-day-old WT mice were inoculated i.c. with EV-D68 IL52 and subsequently inoculated i.p. with anti-CD4, anti-CD8, or isotype control antibody. Mice received a second dose of antibody at 3 dpi. Mice were monitored daily for signs of disease and euthanized upon detection of paralysis. (**A**) Experimental workflow. (**B**) Percent survival (day of paralysis onset) following treatment with either isotype control antibody or anti-CD4 or anti-CD8 antibodies. N = 14-18 mice per group. Data are representative of 2 independent experiments. Log-rank test: *, *P* ≤ 0.05; ****, *P* ≤ 0.0001.

## Discussion

In this study, we used newborn mice to examine the influence of host immunity on replication of EV-D68 in the CNS and development of paralysis. Infection with a neurotropic strain of EV-D68 led to higher levels of cytokines in spinal tissue than did infection with a non-neurotropic strain. These cytokines promoted recruitment of inflammatory cell types to the spinal cord, which is considered an immune-privileged tissue (28). Mice lacking functional immune cell recruitment or functional lymphocytes had diminished disease relative to immunocompetent mice. These data indicate that immune cells contribute to development of EV-D68-associated paralysis in newborn mice, which enhances an understanding of enterovirus neuropathogenesis and provides insights into mechanisms by which host immunity contributes to disease.

Different strains of EV-D68 produced little to no viral load in the brain at 3 dpi in WT mice (Figure 1A), which was the site of inoculation. Failure to replicate efficiently in the brain might be attributable to limited expression of proviral host factors, such as receptors or entry mediators, or a restrictive innate immune response, as mice lacking type I interferon receptors allow more efficient replication of EV-D68 in the brain (25). EV-D68 IL52 was the only strain tested that produced detectable viral loads in spinal tissue of WT mice at 3 dpi (Figure 1A), and the titers of this strain increased in spinal tissue over time (Figure 1B). While *Ccr2^-/-^* and *Rag1^-/-^* mice were less susceptible to EV-D68-mediated disease relative to WT mice, viral loads in spinal tissue of WT and knockout mice at 3 dpi were comparable. These findings suggest that viral replication is necessary, but not sufficient, for development of paralysis in newborn mice.

Inflammation contributes to host immunity and can limit viral replication and disease, but it may augment disease if unregulated or occurs in an immune-privileged site (29). Compared with mice inoculated with avirulent strain MO49 or PBS, mice inoculated with IL52 elaborated higher levels of chemoattractant cytokines CCL2, CCL7, and CCL12 (Figure 2), which mediate immune cell recruitment by binding to CCR2 (30). CCR2 is expressed by several cell types, including dendritic cells, macrophages, monocytes, B cells, T cells, and other immune and non-immune cells (27,30).

Significantly greater numbers of CD8^+^ T cells and macrophages were present in the spinal cord of mice inoculated with IL52 but not MO49 or PBS (Figure 4). This robust immune cell infiltration elicited by IL52 is similar to that observed in post-mortem studies of a child who had flaccid paralysis, in which macrophages, CD8^+^ T cells, and perforin, but not caspase-3, were detected in spinal cord sections (3,8). These observations suggest that the immune response, and not virus-induced apoptosis, contributes to EV-D68 neuropathogenesis.

Mature lymphocytes can impair recovery following spinal cord injury and promote damage to myelin (17). Since antibody-mediated depletion of CD8^+^ T cells significantly reduced paralysis in IL52-infected mice, we conclude that immune-mediated cytotoxicity contributes to motor neuron injury and development of paralysis. T cells are recruited to the spinal cord in a CCR2-dependent manner, as lack of CCR2 results in diminished T cell recruitment during EV-D68 infection (Figure 6). MHC class I molecules are expressed by neurons early during mid-gestation (E9.5-10.5) (31–33). EV-D68-infected neurons marked by presentation of virus-derived peptides in complex with MHC class I molecules could be recognized by CD8^+^ T cells entering spinal tissue and subsequently killed by production of cytolytic molecules such as perforin and granzyme B (34).

Three-day-old WT mice inoculated with strain IL52 developed paralysis by 5-6 dpi and had significantly greater numbers of T cells in spinal tissue than those inoculated with strain MO49 or PBS (Figure 4), suggesting a rapid T-cell response to EV-D68 despite the young age of the mice. Functions of T cells in newborn and adult mice differ (35–37), which may explain this early T cell response to EV-D68 in our experiments. In humans, neonatal CD8^+^ T cells have a distinct pattern of gene expression relative to adult CD8^+^ T cells (38). In newborn mice, naive CD8^+^ T cells are more functionally reactive and rapidly proliferate and become terminally differentiated (39). In addition to MHC class I-restricted recognition of EV-D68 peptides, newborn CD8^+^ T cells might manifest nonspecific innate-like immune functions that lead to neuronal cell death (35,38). Thus, following recruitment of CD8^+^ T cells to the spinal cord of EV-D68-infected newborn mice, neurons could be killed following antigen-restricted cytolysis or a nonspecific innate-like response.

Antibody-mediated depletion of T cells provided substantial but not complete protection against EV-D68-mediated disease, suggesting that additional immune cell subsets contribute to development of paralysis. We think it possible that a multicellular inflammatory environment leads to EV-D68-associated AFM. Other inflammatory immune cells, such as CD86^high^/iNOS^high^ M1 macrophages and Ly6C^high^ monocytes, which express CCR2 and were detected in spinal tissue of paralyzed mice, could influence disease by directly or indirectly injuring neurons (17,40,41). Understanding the contribution of immune cells and their mechanisms of neuronal injury may aid in identifying more specific immune-related therapies for AFM than those used thus far.

Current treatments for EV-D68-mediated AFM are limited. There is insufficient evidence to support the use or avoidance of any specific therapy (42). Treatment of EV-D68-infected mice with dexamethasone, a corticosteroid, was associated with significantly higher loads of EV-D68 and enhanced disease (9). Fluoxetine, an FDA-approved antidepressant, reduced EV-D68 replication in cell culture (43) but had no effect on viral loads in the spinal cord or EV-D68-induced disease in mice (9). Human intravenous immunoglobulin containing EV-D68-neutralizing antibodies reduced viral loads and disease in mice, but only if administered early in the disease course (9).

Other antiviral medications, including pleconaril, pocapavir, and vapendavir, lack significant activity against contemporary strains of EV-D68 at clinically relevant concentrations (44). Although vaccination has been a successful strategy for prevention of poliomyelitis, the large number of potentially paralytogenic enteroviruses make development of vaccines to prevent AFM a challenging undertaking (45). Since administration of T-cell-specific antibodies significantly protected mice against EV-D68-associated paralysis, therapeutics targeting T-cell recruitment or function may be an efficient strategy to treat EV-D68 neurologic disease in humans.

Our studies show that inoculation of newborn mice with a neurotropic strain of EV-D68 results in higher levels of chemoattractant cytokines in spinal tissues. These cytokines promote immune cell recruitment to the spinal cord and, in turn, these immune cells are responsible for paralysis. Administration of antibodies specific for either CD4^+^ or CD8^+^ T cells diminish development of EV-D68-mediated paralysis in newborn mice, with a greater effect observed with CD8^+^ depletion. There are currently no FDA-approved virus-specific therapeutics or vaccines to prevent AFM (43). Our findings provide important insights about the influence of host immunity on EV-D68 replication and pathogenesis in the CNS and illuminate new targets for therapeutic intervention.

## Methods

### Cells and viruses

Rhabdomyosarcoma (RD) cells were propagated at 37°C and 5% CO_2_ in Dulbecco’s modified Eagle’s medium (DMEM) supplemented to contain 10% fetal bovine serum, 100 U/mL penicillin, and 100 μg/mL streptomycin. Infectious cDNAs of US/MO-14-18949 (MO49) and US/IL/14-18952 (IL52) were provided by Dr. Raul Andino (UCSF). The following reagents were obtained through BEI Resources, NIAID, NIH: Plasmid pUC19 containing cDNA from enterovirus D68, USA/Fermon infectious clone EV-D68-R-Fermon (Cat. No. NR 52375). Studies with EV-D68 were conducted in BSL2 containment.

Virus stocks were prepared using RD cells at 33°C by co-transfection of the T7-promoter-containing infectious cDNA plasmids and plasmid expressing T7 RNA polymerase (46). T7opt in pCAGGS was a gift from Benhur Lee (Addgene plasmid # 65974 ; http://n2t.net/addgene:65974 ; RRID:Addgene_65974). When cytopathic effect (CPE) was evident (3-5 dpi), cells were frozen and thawed three times, and lysates were adsorbed to RD cells for 24 hours or until CPE was observed. Cells were collected by scraping, resuspended in a small volume of PBS, frozen and thawed three times, and clarified by centrifugation to prepare high-titer viral stocks.

### Viral plaque assays

Viral titers in cell-culture lysate stocks and tissue homogenates were determined by plaque assay using RD cells maintained at 33°C. Serial 10-fold dilutions of samples in sterile PBS containing calcium and magnesium were adsorbed to RD cells at room temperature (RT) for 1 hour. Monolayers were overlaid with a 1:1 (v/v) mixture of 2% agarose (Invitrogen, 16500500) and 2X MEM (GIBCO) supplemented to contain 20% fetal bovine serum, 200 U/mL penicillin, and 200 μg/mL streptomycin and incubated at 33°C for 48 hours. Plaques were visualized and enumerated following staining with neutral red.

### Mouse experiments

All animal husbandry and experimental procedures were conducted in accordance with U.S. Public Health Service policy and approved by the Institutional Animal Care and Use Committee at the University of Pittsburgh. Mouse experiments were conducted in animal biosafety 2 facilities. Three-day-old mice were inoculated i.c. in the right cerebral hemisphere with 5 µl containing 10^5^ PFU of EV-D68 diluted in sterile PBS using a 30-gauge needle and syringe (Hamilton Company). For survival experiments, mice were monitored daily for signs of disease. Death was not used as an endpoint; mice that had signs of paralysis (e.g., limb dragging and unresponsiveness to stimuli) were euthanized immediately with isoflurane.

For viral replication experiments, mice were euthanized at various intervals post-inoculation, and tissues were collected, weighed, and suspended in 1 mL of PBS. Tissues were homogenized by mechanical disruption with stainless-steel beads using a TissueLyser (QIAGEN) and frozen and thawed three times. Viral titers in tissue homogenates were determined by plaque assay using RD cells. Titers were normalized to the weight of each tissue.

For antibody depletion experiments, 3-day-old mice were inoculated i.c. as described and administered 50 µg of anti-keyhole limpet hemocyanin IgG2b isotype (Bio X Cell, BE0090), anti-CD4 clone GK1.5 (Bio X Cell, BE0003-1), or anti-CD8 clone 2.43 (Bio X Cell, BE0061) intraperitoneally in a volume of 30 µl of sterile *InVivo*Pure dilution buffers recommended for each clone (Bio X Cell). Mice were administered a second antibody dose 3 days after the first and were monitored for signs of disease.

### Serum neutralization

Blood was obtained from euthanized mice and allowed to coagulate at RT for 30 minutes. Sera were obtained by clarifying coagulated blood and heat-inactivated at 56°C for 30 minutes. Sera were serially diluted 1:2 (v/v) in completed DMEM, and virus was diluted to a final concentration of 100 TCID_50_ per well. Virus was incubated with serum dilutions at 33°C for 1 hour and inoculated onto RD cells. At 5 dpi, cells were fixed and stained with crystal violet.

### Human spinal cord organoid cultivation and infection

Human iPSCs (SCTi003-A, STEMCELL Technologies, 200-0511) were maintained in mTeSR Plus pluripotent stem cell medium (STEMCELL Technologies, 100-0276) supplemented to contain 10 µM Y-27632 (Tocris, 1254) in flasks coated with 150 µg/mL Cultrex (R&D Systems, 3434-005-02). To prepare 3-DiSC hSCOs, a single-cell suspension of iPSCs was prepared using ACCUMAX (STEMCELL Technologies), and cells were seeded at a density of 9,000 cells/well in 96-well, round-bottom, low-adhesion plates in 100 µL of N2B27 differentiation medium (1:1 [v/v] of DMEM/F-12 [GIBCO] and neurobasal medium [GIBCO] supplemented to contain 10% Knockout serum replacement [Invitrogen], 0.5% N2 supplement [Thermo Fisher Scientific], and 1% B27 supplement without vitamin A [Invitrogen] supplemented to contain 1 mM L-glutamine [Gibco], 0.1 mM 2-mercaptoethanol [Sigma], and 0.5 µM ascorbic acid [Sigma]) as described (11). Every 3 days during differentiation, 50% of the medium was replaced with fresh medium. For patterning, on days 0-3, the medium was supplemented to contain 10 µM Y-27632 (Tocris, 1254), 20 ng/mL bFGF (Thermo Fisher Scientific, 13256-029), and 3 µM CHIR 99021 (Tocris, 4423). From days 0 to 6, the medium was supplemented to contain 10 µM SB431542 (Tocris, 1614). From days 3 to 15, the medium was supplemented to contain 100 nM retinoic acid (Tocris, 302-79-4) and 500 nM smoothened agonist (SAG, Tocris, 912545-86-9).

For infection experiments, hSCOs were plated in pools of 8-12 organoids per well and inoculated with PBS or virus (10^5^ PFU/pool) at RT for 1 h. Organoids were washed three times with PBS and transferred to a new well prior to incubation with fresh medium at 33°C for the duration of the experiment.

### Immunofluorescence

Antibody staining of hSCOs was conducted using the fructose-glycerol clearing method as described (47). hSCOs were fixed with 4% PFA and washed with PBS and 0.1% Tween-20 (v/v) supplemented to contain 0.2% (m/v) bovine serum albumin (BSA) at 4°C. hSCOs subsequently were washed three times with organoid wash buffer (0.1% Triton-X-100 [v/v] and 0.2% BSA [m/v] in PBS). hSCOs were incubated with VP1-specific antibody (Genetex, GTX132313) at a 1:250 dilution, goat anti-rabbit IgG (Thermo Fisher Scientific, A-11008) at a 1:1000 dilution, and phalloidin (Biotium, 00043) overnight at 4°C. hSCOs were cleared with 60% glycerol in 2.5M fructose for 30 min, mounted on slides, and imaged using a Leica Stellaris 5 confocal microscope. Images were processed using Fiji (48).

Mice were inoculated with either PBS or EV-D68 IL52 and euthanized when paralysis was first detected in the IL52-inoculated animals. Spinal columns were dissected and fixed with 10% formalin at RT for 3 days. Spinal cords were dissected and placed in SpineRacks (49), covered with optimal cutting temperature OCT compound, and frozen. Frozen sections were processed and immunostained as described (50). Sections were fixed with 4% formaldehyde at RT for 15 minutes, rinsed three times with PBS, permeabilized with 0.1% Triton X-100 in PBS for 10 min, rinsed three times with PBS, and blocked overnight at 4°C in PBS supplemented to contain 5% BSA. Serial sections were stained with VP1-specific antibody (Genetex, GTX132313) at a 1:250 dilution or NeuN (BioLegend, 834501) at RT for 1 h, rinsed three times with PBS, and stained with goat anti-rabbit IgG (Thermo Fisher Scientific, A-11035) and goat anti-mouse IgG (Thermo Fisher Scientific, A-11001) both at a 1:1000 dilution at RT for 1h. Sections were counterstained with DAPI.

### Luminex assays

Brain or spinal tissue homogenates prepared from samples used for viral titer determination were analyzed by Luminex profiling (Luminex Corporation) with the Bio-Plex Pro Mouse Chemokine 31-plex panel (Bio-Rad, 12009159) according to the manufacturer’s instructions. Cytokine levels were quantified using the LabMAP multianalyte profiling system (Luminex).

Supernatants were collected from hSCOs inoculated with PBS or EV-D68 at 3 dpi, and cytokine levels were determined with the Bio-Plex Human Inflammation Panel 1 37-plex assay kit (Bio-Rad, 171AL001M) according to the manufacturer’s instructions using the MAGPIX laboratory multianalyte profiling system (Millipore) developed by Luminex.

### Flow cytometric analyses of neonatal mouse spinal cord samples

Mice were inoculated with either PBS or EV-D68 IL52 and euthanized when paralysis was first detected in the IL52-inoculated animals. Spinal cords were resected and submerged in ice-cold Hanks’ balanced salt solution (HBSS) (GIBCO). Spinal cords were incubated with 0.5 mg/ml trypsin (Worthington, LS003708) in HBSS at RT for 30 min, washed twice with HBSS, and manually dissociated with a fire-polished borosilicate glass Pasteur pipette. Cell suspensions were passed through a 70-μm cell strainer (Falcon, 352350).

Cells were collected to form a pellet by centrifugation at 1500 rpm for 4 minutes, resuspended in FACS buffer, and incubated with the antibodies and stains shown in Supplemental Table 1. Single-cell suspensions were washed with PBS and stained with LIVE/DEAD Fixable Violet dye (Thermo Fisher Scientific, L34963) 1:1000 in PBS for 15 minutes. Cells were washed with FACS buffer (2% heat-inactivated FBS in PBS) and blocked with CD16/CD32 specific antibody (Tonbo Biosciences, 70-0161-M001) and True-stain monocytes blocker (BioLegend, Inc., 46102) according to the manufacturer’s protocol. Cells were stained with antibodies specific for surface proteins (Supplemental Table 1) (1:100, v/v) in Brilliant Stain buffer (BD, 566349) at 4°C for 30 minutes and washed with FACS buffer. Cells were fixed and permeabilized using Cytofix/Cytoperm solution (BD, 554714) at RT for 30 minutes. Cells were stained with antibodies specific for intracellular proteins (Supplemental Table 1) (1:100,v/v) in Perm/Wash buffer (BD) supplemented to contain 2% rat serum at RT for 1 hour. Cells were washed with Perm/Wash buffer, resuspended in FACS buffer, and analyzed using an Aurora multispectral flow cytometer (Cytek). Absolute cell number was determined by comparing to Precision Count Beads (BioLegend, 424902). Cell populations were identified using sequential gating strategy as described (51,52).

### Statistical analysis

The difference between groups were assessed using unpaired two-tailed Student’s *t* tests or log-rank tests. Error bars in figures represent standard errors of the mean. A *P* value of < 0.05 was considered statistically significant. All analyses of data were conducted using GraphPad Prism (version 10.1.2).

## Author contributions

MAWA, JL, JVW, MCF, and TSD designed research studies. MAWA, JL, SM, JEJ, and MCF conducted experiments. MAWA, JL, SM, JEJ, and MCF acquired data. MAWA, JL, SM, JEJ, JVW, MCF, and TSD analyzed data. MAWA, MCF, and TSD wrote the manuscript.

## Acknowledgments

We thank members of the Dermody, Freeman, and Williams lab for essential discussions and suggestions during the conduct of this research. We are grateful to Dr. Taylor Eddens for advice about the T-cell depletion studies, Dr. Jorna Sojati for technical assistance with the Luminex assays, and Paul Culler for 3D printing Spine Racks. The confocal images were captured in the Cell Imaging Core Facility at UPMC Children’s Hospital of Pittsburgh Rangos Research Center. We are grateful to the Cell Imaging Core staff for technical assistance. We thank Drs. Pengcheng Shang and Danica Sutherland for careful review of the manuscript. The graphical abstract accompanying this manuscript was prepared using BioRender. Woods Acevedo, M. (2024) BioRender.com/ k97m275.

This work was funded by NIH T32 AI049820 (M.A.W.A), a Helen Hay Whitney Foundation Fellowship (M.A.W.A.), a Burroughs Wellcome Fund Postdoctoral Enrichment Program award (M.A.W.A.), NIH K08 AI171177 (M.C.F.), and the Richard King Mellon Foundation (M.A.W.A., J.V.W., M.C.F., and T.S.D.). Additional funding was provided by the Henry L. Hillman Foundation (J.V.W.) and the Heinz Endowments (T.S.D.). The funders had no role in study design, data collection and interpretation, or the decision to submit the work for publication.

## Supplemental Figure Legends

**Supplemental Figure 1.**
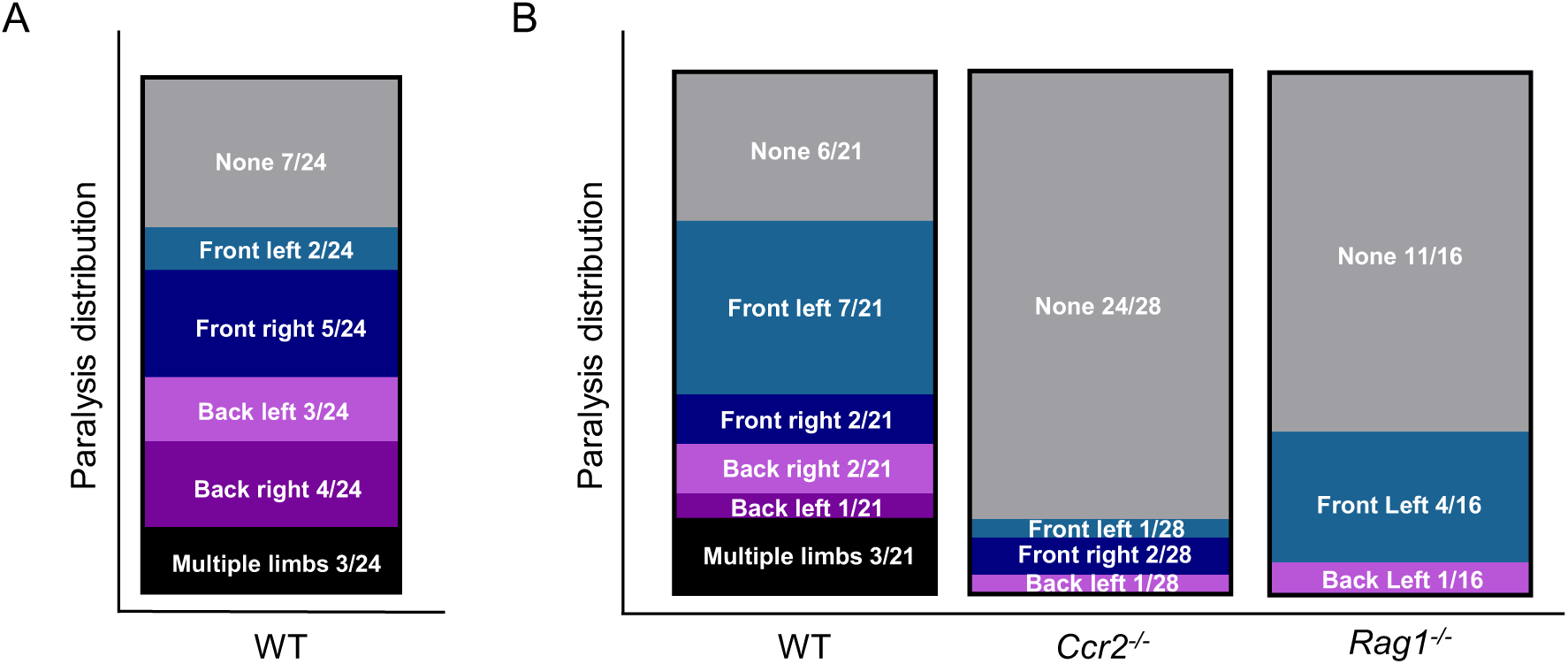
Paralyzed limb distribution of US/IL/14-18952-inoculated mice. Three-day-old mice of the indicated genotypes were inoculated i.c. with 10^5^ PFU of EV-D68 IL52, monitored daily for disease, and euthanized upon signs of paralysis. Distribution of limb paralysis from experiments presented in (**A**) Figure 1D and (**B**) Figures 5B and 7B. Experiments shown in Figures 5B and 7B were conducted concurrently. Data are representative of 2-3 independent experiments.

**Supplemental Table 1.**
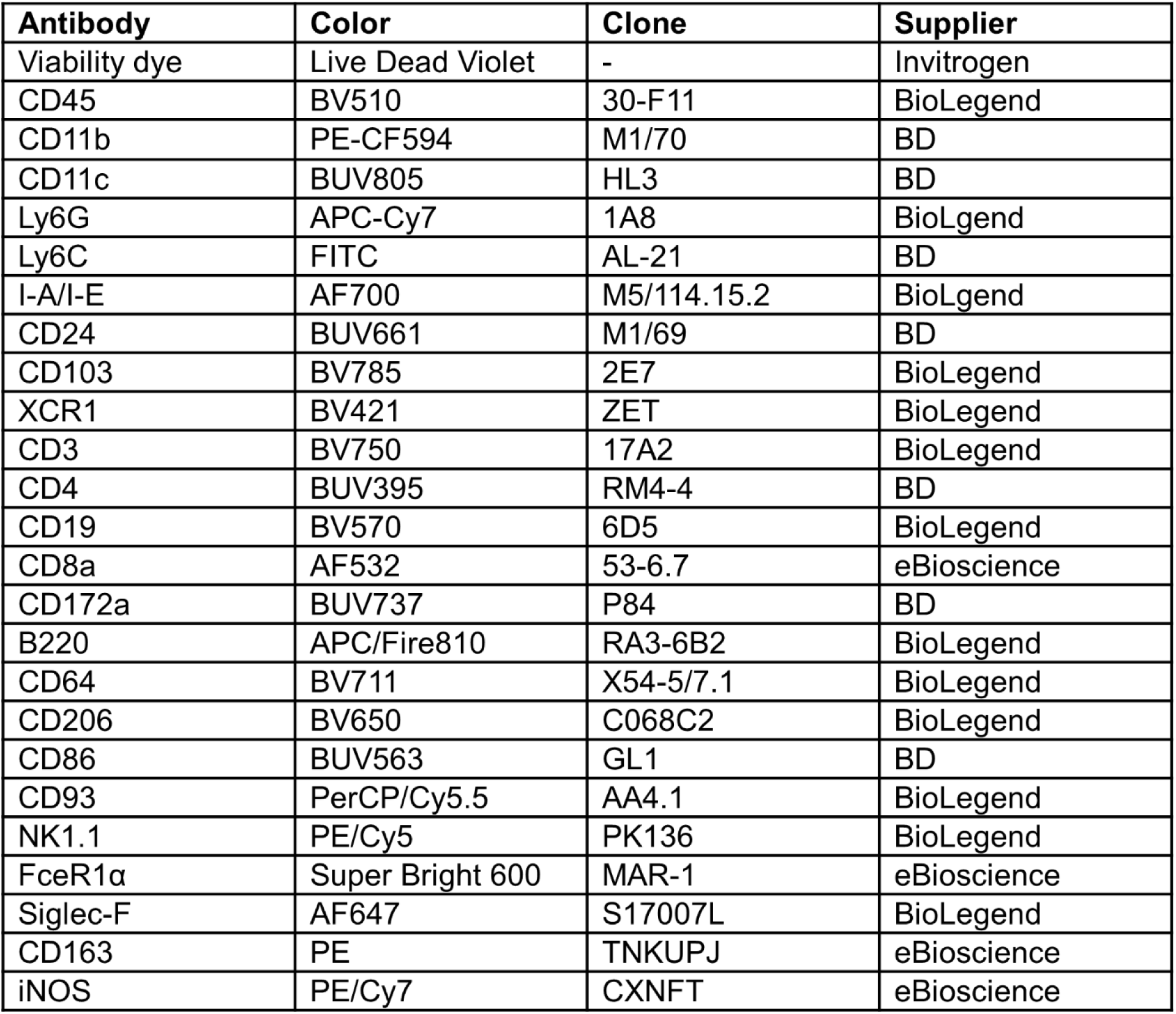
Antibody and dye reagents used for flow cytometry analysis.

| **Antibody** | **Color** | **Clone** | **Supplier** |
| --- | --- | --- | --- |
| Viability dye | Live Dead Violet | - | Invitrogen |
| CD45 | BV510 | 30-F11 | BioLegend |
| CD11b | PE-CF594 | M1/70 | BD |
| CD11c | BUV805 | HL3 | BD |
| Ly6G | APC-Cy7 | 1A8 | BioLegend |
| Ly6C | FITC | AL-21 | BD |
| I-A/I-E | AF700 | M5/114.15.2 | BioLegend |
| CD24 | BUV661 | M1/69 | BD |
| CD103 | BV785 | 2E7 | BioLegend |
| XCR1 | BV421 | ZET | BioLegend |
| CD3 | BV750 | 17A2 | BioLegend |
| CD4 | BUV395 | RM4-4 | BD |
| CD19 | BV570 | 6D5 | BioLegend |
| CD8a | AF532 | 53-6.7 | eBioscience |
| CD172a | BUV737 | P84 | BD |
| B220 | APC/Fire810 | RA3-6B2 | BioLegend |
| CD64 | BV711 | X54-5/7.1 | BioLegend |
| CD206 | BV650 | C068C2 | BioLegend |
| CD86 | BUV563 | GL1 | BD |
| CD93 | PerCP/Cy5.5 | AA4.1 | BioLegend |
| NK1.1 | PE/Cy5 | PK136 | BioLegend |
| FcεR1α | Super Bright 600 | MAR-1 | eBioscience |
| Siglec-F | AF647 | S17007L | BioLegend |
| CD163 | PE | TNKUPJ | eBioscience |
| iNOS | PE/Cy7 | CXNFT | eBioscience |

**Supplemental Figure 2.**
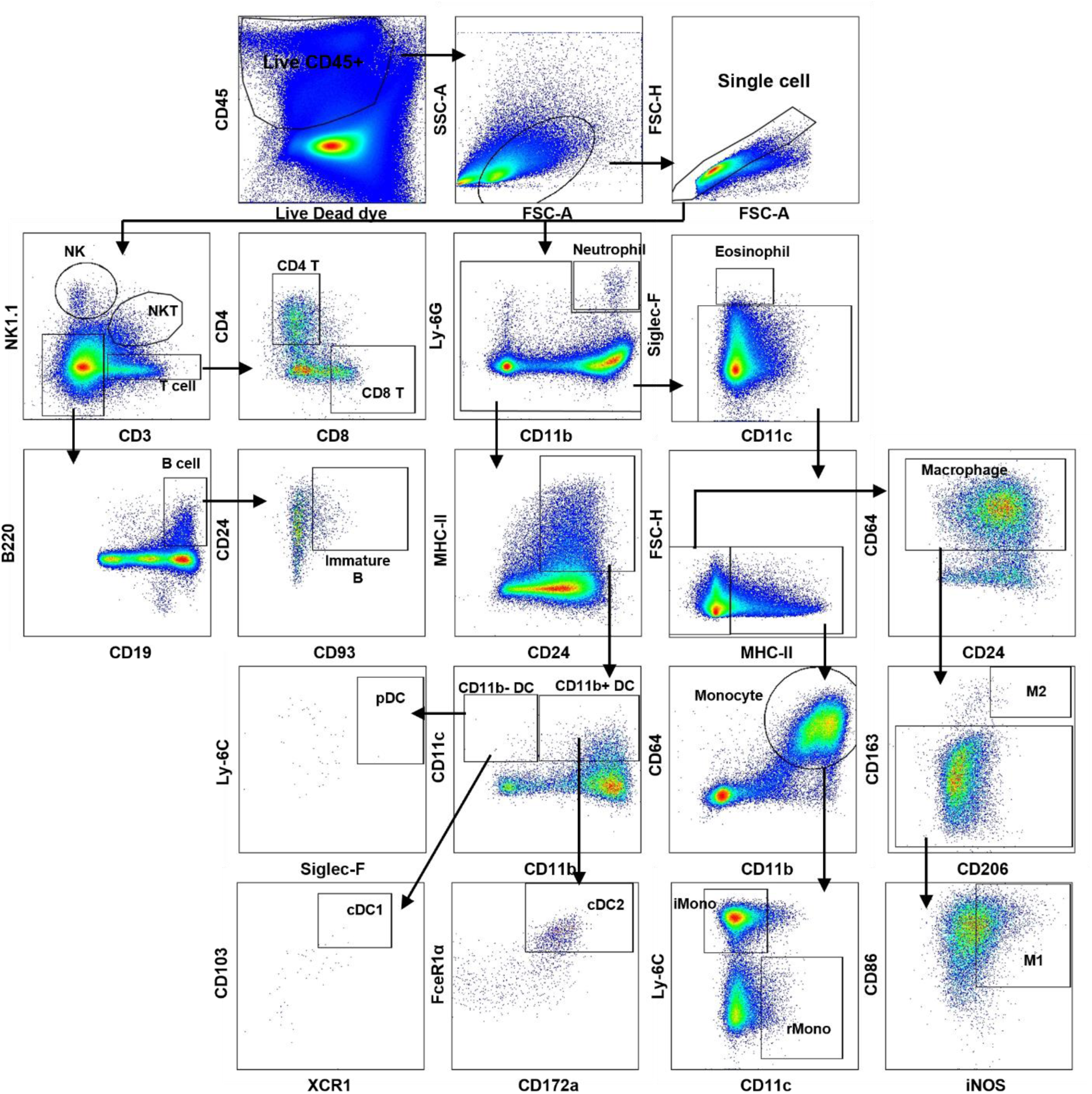
Gating strategy used to identify immune cell populations in the spinal cord. Flow cytometric gating strategy for mouse leukocytes in the spinal cord. Frequency and absolute cell numbers of different subpopulations of immune cells were assessed. Representative pseudocolored dot density plots from EV-D68 IL52-inoculated WT mice. Boxed or circled populations labelled with cell-type-specific antibodies indicate populations of interest.

**Supplemental Figure 3.**
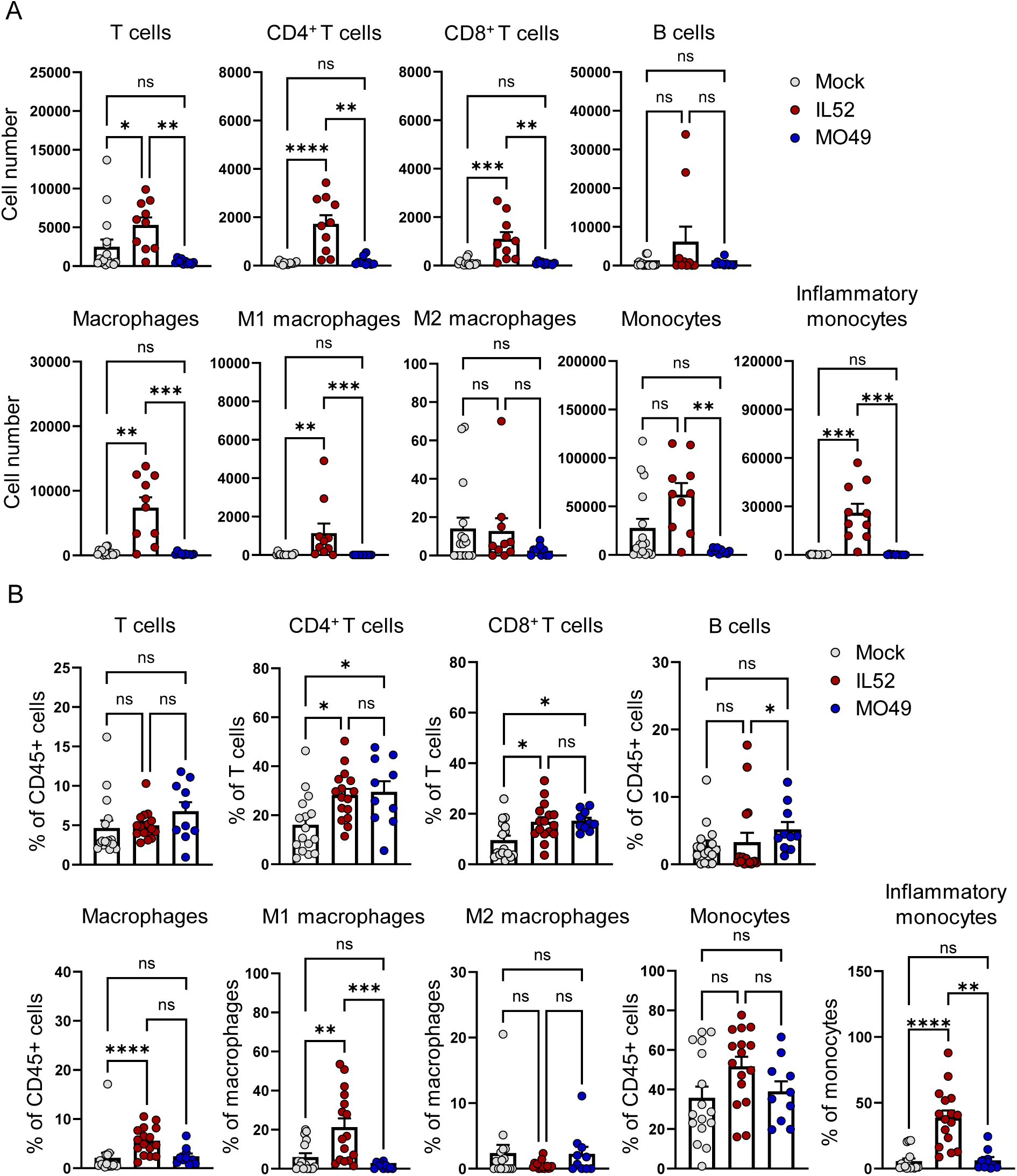
Mice inoculated with neurovirulent EV-D68 have altered populations of immune cells in the spinal cord. Three-day-old WT mice were inoculated i.c. with PBS (mock) or EV-D68 MO49 or IL52. Spinal cords were resected from paralyzed mice inoculated with IL52 or day-matched mice inoculated with MO49 or PBS. Single-cell suspensions were prepared, stained, and analyzed by flow cytometry. (**A**) Numbers and (**B**) percentages of selected cell types are shown. Data are representative of 2-4 independent experiments. Each symbol represents an individual mouse. Error bars indicate mean ± SEM. Mann-Whitney test: *, *P* ≤ 0.05; **, *P* ≤ 0.01; ***, *P* ≤ 0.001; ****, *P* ≤ 0.0001; ns = not significant.

**Supplemental Figure 4.**
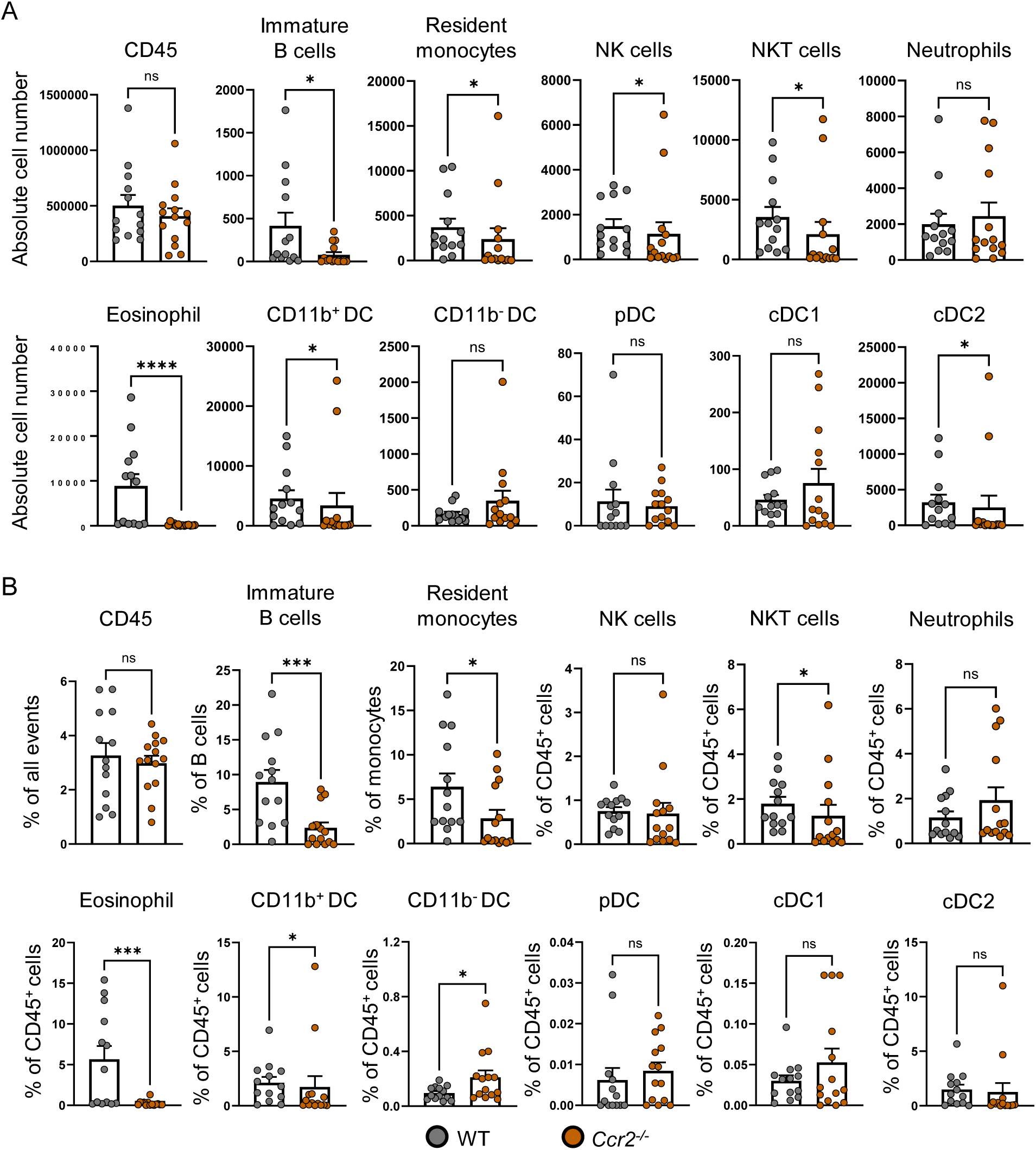
C*c*r2*^-/-^* mice have altered immune cell recruitment following neurovirulent EV-D68 inoculation. Three-day-old WT or *Ccr2^-/-^* mice were inoculated i.c. with EV-D68 IL52. Spinal cords were resected from paralyzed WT mice or day-matched *Ccr2^-/-^* mice. Single-cell suspensions were prepared, stained, and analyzed by flow cytometry. (**A**) Numbers and (**B**) percentages of selected cell types are shown. Each symbol represents an individual mouse. Error bars indicate mean ± SEM. Mann-Whitney test: *, *P* ≤ 0.05; **, *P* ≤ 0.01; ***, *P* ≤ 0.001; ****, *P* ≤ 0.0001; ns = not significant.

**Supplemental Figure 5.**
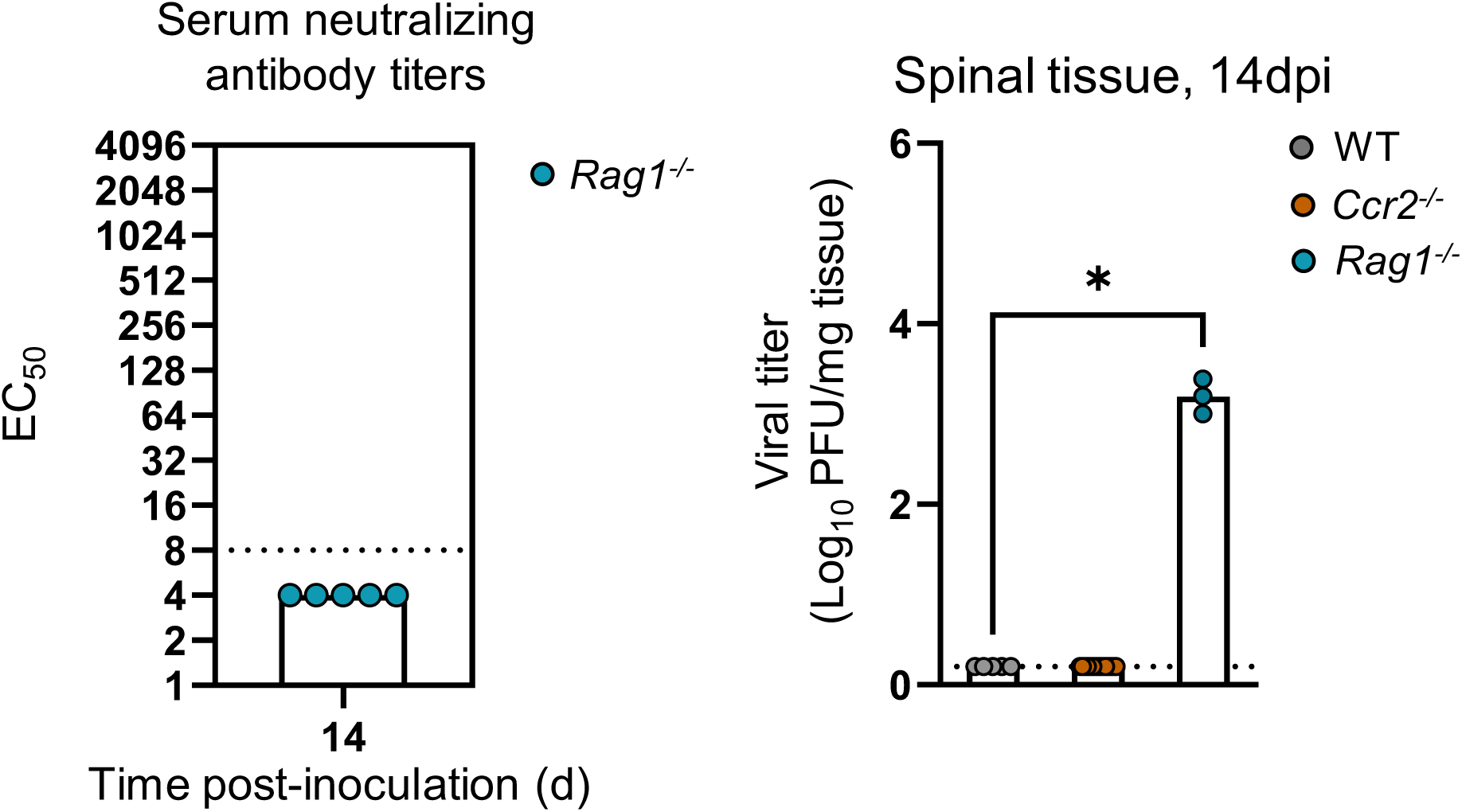
Neurovirulent EV-D68-inoculated *Rag1^-/-^* mice fail to produce neutralizing antibodies or clear virus. Three-day-old WT, *Ccr2^-/-^*, or *Rag1^-/-^* mice were inoculated i.c. with EV-D68 IL52. (**A**) Serum was collected at 14 dpi from IL52-inoculated *Rag1^-/-^* mice and assessed for neutralizing antibody titers. Dotted line indicates the limit of detection. (**B**) Spinal tissue was resected at 14 dpi, and viral titers were determined by plaque assay. Each symbol represents an individual mouse. Mann-Whitney test: *, *P* ≤ 0.05.

**Supplemental Figure 6.**
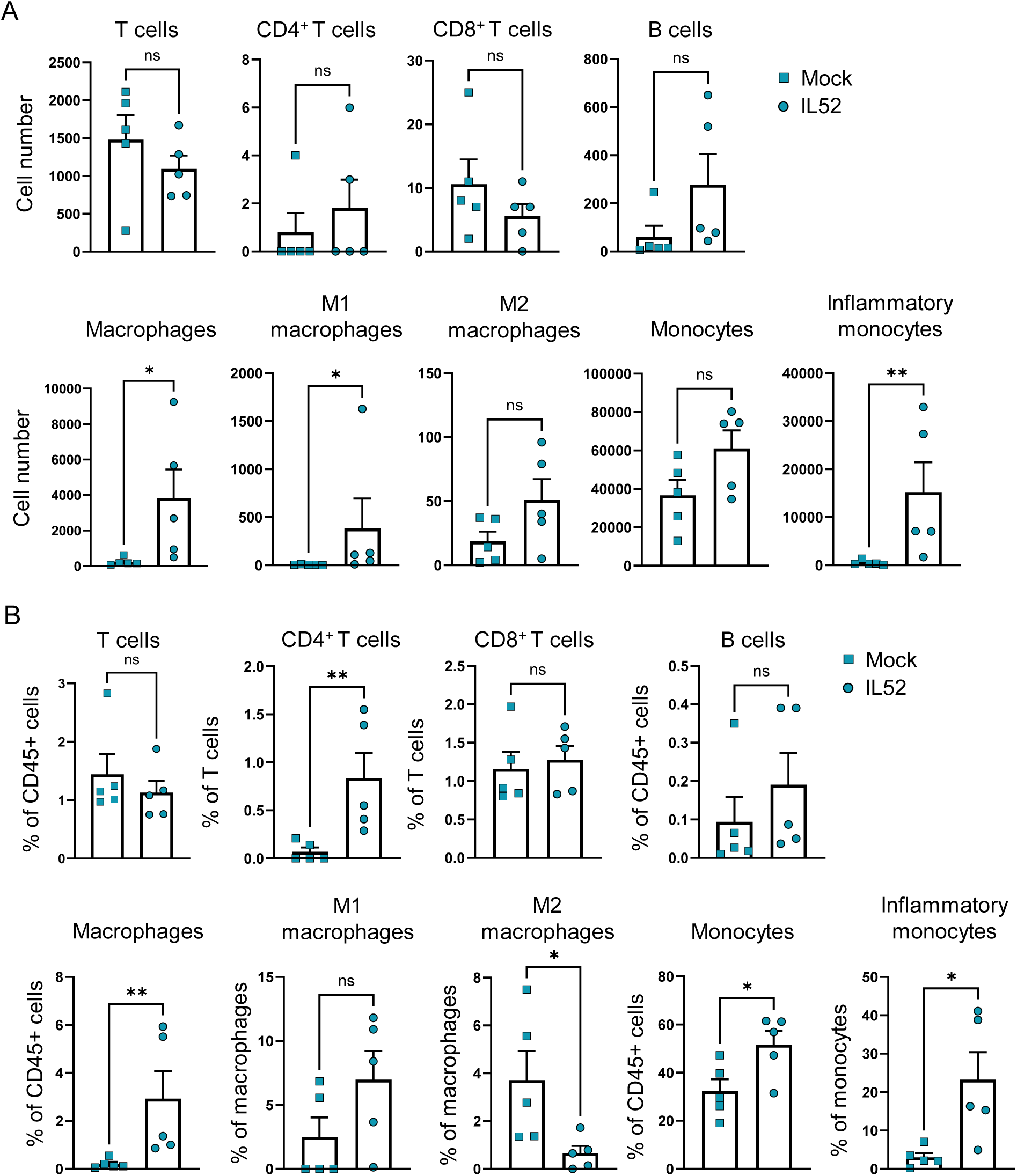
R*a*g1*^-/-^* mice have varied macrophage and T-cell populations following neurovirulent EV-D68 inoculation. Three-day-old *Rag1^-/-^* mice were inoculated i.c. with PBS (mock) or EV-D68 IL52. Spinal cords were resected from paralyzed *Rag1^-/-^* mice or day-matched mock-inoculated mice. Single-cell suspensions were prepared, stained, and analyzed by flow cytometry. (**A**) Numbers and (**B**) percentages of selected cell types are shown. Data are representative of 2-3 independent experiments. Each symbol represents an individual mouse. Error bars indicate mean ± SEM. Mann-Whitney test: *, *P* ≤ 0.05; **, *P* ≤ 0.01; ns = not significant.

**Supplemental Figure 7.**
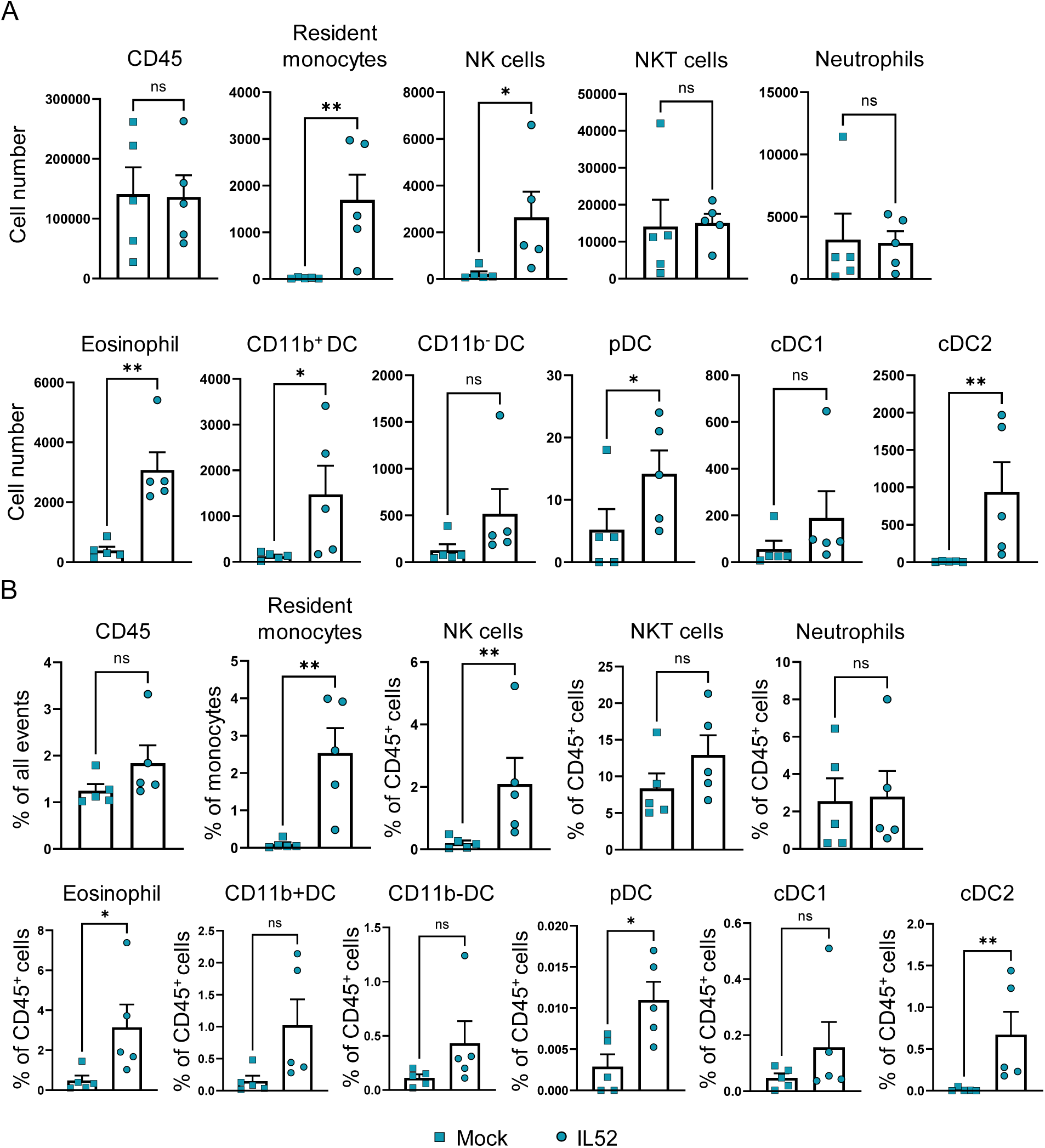
Mice lacking mature lymphocytes have altered immune cell recruitment following neurovirulent EV-D68 inoculation. Three-day-old *Rag1^-/-^* mice were inoculated i.c. with EV-D68 IL52. Spinal cords were resected from paralyzed *Rag1^-/-^* mice or day-matched mock-inoculated mice. Single-cell suspensions were prepared, stained, and analyzed by flow cytometry. (**A**) Numbers and (**B**) percentages of selected cell types are shown. Error bars are mean ± SEM. *, *P* ≤ 0.05; **, *P* ≤ 0.01; ns = not significant, Mann-Whitney test. Each symbol represents an individual mouse. Data are representative of 2-3 independent experiments.

## Notes

### Competing Interest Statement

The authors have declared no competing interest.

## References

1. Brown BA, Nix WA, Sheth M, Frace M, Oberste MS. Seven Strains of Enterovirus D68 Detected in the United States during the 2014 Severe Respiratory Disease Outbreak. Genome Announc. Nov 20 2014;2(6)doi:10.1128/genomeA.01201-14

2. Midgley CM, Watson JT, Nix WA, et al. Severe respiratory illness associated with a nationwide outbreak of enterovirus D68 in the USA (2014): a descriptive epidemiological investigation. Lancet Respir Med. Nov 2015;3(11):879–87. doi:10.1016/S2213-2600(15)00335-5

3. Kreuter JD, Barnes A, McCarthy JE, et al. A fatal central nervous system enterovirus 68 infection. Arch Pathol Lab Med. Jun 2011;135(6):793–6. doi:10.5858/2010-0174-CR.1

4. Hixon AM, Yu G, Leser JS, et al. A mouse model of paralytic myelitis caused by enterovirus D68. PLoS Pathog. Feb 2017;13(2):e1006199. doi:10.1371/journal.ppat.1006199

5. Schieble JH, Fox VL, Lennette EH. A probable new human picornavirus associated with respiratory diseases. Am J Epidemiol. Mar 1967;85(2):297–310. doi:10.1093/oxfordjournals.aje.a120693

6. Sejvar JJ, Lopez AS, Cortese MM, et al. Acute Flaccid Myelitis in the United States, August-December 2014: Results of Nationwide Surveillance. Clin Infect Dis. Sep 15 2016;63(6):737–745. doi:10.1093/cid/ciw372

7. Shah MM, Perez A, Lively JY, et al. Enterovirus D68-Associated Acute Respiratory Illness horizontal line New Vaccine Surveillance Network, United States, July-November 2018-2020. MMWR Morb Mortal Wkly Rep. Nov 26 2021;70(47):1623–1628. doi:10.15585/mmwr.mm7047a1

8. Vogt MR, Wright PF, Hickey WF, De Buysscher T, Boyd KL, Crowe JE, Jr. Enterovirus D68 in the Anterior Horn Cells of a Child with Acute Flaccid Myelitis. N Engl J Med. May 26 2022;386(21):2059–2060. doi:10.1056/NEJMc2118155

9. Hixon AM, Clarke P, Tyler KL. Evaluating Treatment Efficacy in a Mouse Model of Enterovirus D68-Associated Paralytic Myelitis. J Infect Dis. Dec 5 2017;216(10):1245–1253. doi:10.1093/infdis/jix468

10. Girard S, Couderc T, Destombes J, Thiesson D, Delpeyroux F, Blondel B. Poliovirus induces apoptosis in the mouse central nervous system. J Virol. Jul 1999;73(7):6066–72. doi:10.1128/JVI.73.7.6066-6072.1999

11. Aguglia G, Coyne CB, Dermody TS, Williams JV, Freeman MC. Contemporary enterovirus-D68 isolates infect human spinal cord organoids. mBio. Aug 31 2023;14(4):e0105823. doi:10.1128/mbio.01058-23

12. Lopez A, Lee A, Guo A, et al. Vital Signs: Surveillance for Acute Flaccid Myelitis - United States, 2018. MMWR Morb Mortal Wkly Rep. Jul 12 2019;68(27):608–614. doi:10.15585/mmwr.mm6827e1

13. Mishra N, Ng TFF, Marine RL, et al. Antibodies to Enteroviruses in Cerebrospinal Fluid of Patients with Acute Flaccid Myelitis. mBio. Aug 13 2019;10(4)doi:10.1128/mBio.01903-19

14. Schubert RD, Hawes IA, Ramachandran PS, et al. Pan-viral serology implicates enteroviruses in acute flaccid myelitis. Nat Med. Nov 2019;25(11):1748–1752. doi:10.1038/s41591-019-0613-1

15. Jurado KA, Yockey LJ, Wong PW, Lee S, Huttner AJ, Iwasaki A. Antiviral CD8 T cells induce Zika-virus-associated paralysis in mice. Nat Microbiol. Feb 2018;3(2):141–147. doi:10.1038/s41564-017-0060-z

16. Kenney LL, Carter EP, Gil A, Selin LK. T cells in the brain enhance neonatal mortality during peripheral LCMV infection. PLoS Pathog. Jan 2021;17(1):e1009066. doi:10.1371/journal.ppat.1009066

17. Wu B, Matic D, Djogo N, Szpotowicz E, Schachner M, Jakovcevski I. Improved regeneration after spinal cord injury in mice lacking functional T- and B-lymphocytes. Exp Neurol. Oct 2012;237(2):274–85. doi:10.1016/j.expneurol.2012.07.016

18. Vermillion MS, Dearing J, Zhang Y, et al. Animal Models of Enterovirus D68 Infection and Disease. J Virol. Aug 10 2022;96(15):e0083322. doi:10.1128/jvi.00833-22

19. Brown DM, Hixon AM, Oldfield LM, et al. Contemporary Circulating Enterovirus D68 Strains Have Acquired the Capacity for Viral Entry and Replication in Human Neuronal Cells. mBio. Oct 16 2018;9(5)doi:10.1128/mBio.01954-18

20. Sooksawasdi Na Ayudhya S, Meijer A, Bauer L, et al. Enhanced Enterovirus D68 Replication in Neuroblastoma Cells Is Associated with a Cell Culture-Adaptive Amino Acid Substitution in VP1. mSphere. Nov 4 2020;5(6)doi:10.1128/mSphere.00941-20

21. Rosenfeld AB, Warren AL, Racaniello VR. Neurotropism of Enterovirus D68 Isolates Is Independent of Sialic Acid and Is Not a Recently Acquired Phenotype. mBio. Oct 22 2019;10(5)doi:10.1128/mBio.02370-19

22. Poelaert KCK, van Kleef R, Liu M, et al. Enterovirus D-68 Infection of Primary Rat Cortical Neurons: Entry, Replication, and Functional Consequences. mBio. Apr 25 2023;14(2):e0024523. doi:10.1128/mbio.00245-23

23. Laksono BM, Sooksawasdi Na Ayudhya S, Aguilar-Bretones M, Embregts CWE, van Nierop GP, van Riel D. Human B cells and dendritic cells are susceptible and permissive to enterovirus D68 infection. mSphere. Feb 28 2024;9(2):e0052623. doi:10.1128/msphere.00526-23

24. Zhang C, Zhang X, Dai W, et al. A Mouse Model of Enterovirus D68 Infection for Assessment of the Efficacy of Inactivated Vaccine. Viruses. Jan 30 2018;10(2)doi:10.3390/v10020058

25. Yeh MT, Capponi S, Catching A, Bianco S, Andino R. Mapping Attenuation Determinants in Enterovirus-D68. Viruses. Aug 8 2020;12(8)doi:10.3390/v12080867

26. Grizer CS, Messacar K, Mattapallil JJ. Enterovirus-D68 - A Reemerging Non-Polio Enterovirus that Causes Severe Respiratory and Neurological Disease in Children. Front Virol. 2024;4doi:10.3389/fviro.2024.1328457

27. Martin E, Delarasse C. Complex role of chemokine mediators in animal models of Alzheimer’s Disease. Biomed J. Feb 2018;41(1):34–40. doi:10.1016/j.bj.2018.01.002

28. Forrester JV, McMenamin PG, Dando SJ. CNS infection and immune privilege. Nat Rev Neurosci. Nov 2018;19(11):655–671. doi:10.1038/s41583-018-0070-8

29. Chitnis T, Weiner HL. CNS inflammation and neurodegeneration. J Clin Invest. Oct 2 2017;127(10):3577–3587. doi:10.1172/JCI90609

30. She S, Ren L, Chen P, et al. Functional Roles of Chemokine Receptor CCR2 and Its Ligands in Liver Disease. Front Immunol. 2022;13:812431. doi:10.3389/fimmu.2022.812431

31. Chacon MA, Boulanger LM. MHC class I protein is expressed by neurons and neural progenitors in mid-gestation mouse brain. Mol Cell Neurosci. Jan 2013;52:117–27. doi:10.1016/j.mcn.2012.11.004

32. Cebrian C, Loike JD, Sulzer D. Neuronal MHC-I expression and its implications in synaptic function, axonal regeneration and Parkinson’s and other brain diseases. Front Neuroanat. 2014;8:114. doi:10.3389/fnana.2014.00114

33. Lazarczyk MJ, Eyford BA, Varghese M, et al. The intracellular domain of major histocompatibility class-I proteins is essential for maintaining excitatory spine density and synaptic ultrastructure in the brain. Sci Rep. Apr 20 2023;13(1):6448. doi:10.1038/s41598-023-30054-8

34. Sun L, Su Y, Jiao A, Wang X, Zhang B. T cells in health and disease. Signal Transduct Target Ther. Jun 19 2023;8(1):235. doi:10.1038/s41392-023-01471-y

35. Rudd BD. Neonatal T Cells: A Reinterpretation. Annu Rev Immunol. Apr 26 2020;38:229–247. doi:10.1146/annurev-immunol-091319-083608

36. Wissink EM, Smith NL, Spektor R, Rudd BD, Grimson A. MicroRNAs and Their Targets Are Differentially Regulated in Adult and Neonatal Mouse CD8+ T Cells. Genetics. Nov 2015;201(3):1017–30. doi:10.1534/genetics.115.179176

37. Le Campion A, Gagnerault MC, Auffray C, et al. Lymphopenia-induced spontaneous T-cell proliferation as a cofactor for autoimmune disease development. Blood. Aug 27 2009;114(9):1784–93. doi:10.1182/blood-2008-12-192120

38. Galindo-Albarran AO, Lopez-Portales OH, Gutierrez-Reyna DY, et al. CD8(+) T Cells from Human Neonates Are Biased toward an Innate Immune Response. Cell Rep. Nov 15 2016;17(8):2151–2160. doi:10.1016/j.celrep.2016.10.056

39. Smith NL, Wissink E, Wang J, et al. Rapid proliferation and differentiation impairs the development of memory CD8+ T cells in early life. J Immunol. Jul 1 2014;193(1):177–84. doi:10.4049/jimmunol.1400553

40. Li Q, Barres BA. Microglia and macrophages in brain homeostasis and disease. Nat Rev Immunol. Apr 2018;18(4):225–242. doi:10.1038/nri.2017.125

41. Fricker M, Oliva-Martin MJ, Brown GC. Primary phagocytosis of viable neurons by microglia activated with LPS or Abeta is dependent on calreticulin/LRP phagocytic signalling. J Neuroinflammation. Aug 13 2012;9:196. doi:10.1186/1742-2094-9-196

42. Centers for Disease Control and Prevention. Clinical Guidance for the Acute Medical Treatment of AFM. https://www.cdc.gov/acute-flaccid-myelitis/hcp/clinical-guidance/index.html

43. Messacar K, Sillau S, Hopkins SE, et al. Safety, tolerability, and efficacy of fluoxetine as an antiviral for acute flaccid myelitis. Neurology. Apr 30 2019;92(18):e2118–e2126. doi:10.1212/WNL.0000000000006670

44. Rhoden E, Zhang M, Nix WA, Oberste MS. In Vitro Efficacy of Antiviral Compounds against Enterovirus D68. Antimicrob Agents Chemother. Dec 2015;59(12):7779–81. doi:10.1128/AAC.00766-15

45. Moss DL, Paine AC, Krug PW, Kanekiyo M, Ruckwardt TJ. Enterovirus virus-like-particle and inactivated poliovirus vaccines do not elicit substantive cross-reactive antibody responses. PLoS Pathog. Apr 2024;20(4):e1012159. doi:10.1371/journal.ppat.1012159

46. Yun T, Park A, Hill TE, et al. Efficient reverse genetics reveals genetic determinants of budding and fusogenic differences between Nipah and Hendra viruses and enables real-time monitoring of viral spread in small animal models of henipavirus infection. J Virol. Jan 15 2015;89(2):1242–53. doi:10.1128/JVI.02583-14

47. Dekkers JF, Alieva M, Wellens LM, et al. High-resolution 3D imaging of fixed and cleared organoids. Nat Protoc. Jun 2019;14(6):1756–1771. doi:10.1038/s41596-019-0160-8

48. Schindelin J, Arganda-Carreras I, Frise E, et al. Fiji: an open-source platform for biological-image analysis. Nat Methods. Jun 28 2012;9(7):676–82. doi:10.1038/nmeth.2019

49. Fiederling F, Hammond LA, Ng D, Mason C, Dodd J. SpineRacks and SpinalJ for efficient analysis of neurons in a 3D reference atlas of the mouse spinal cord. STAR Protoc. Dec 17 2021;2(4):100897. doi:10.1016/j.xpro.2021.100897

50. Zhou X, Moore BB. Lung Section Staining and Microscopy. Bio Protoc. May 20 2017;7(10)doi:10.21769/BioProtoc.2286

51. Misharin AV, Morales-Nebreda L, Mutlu GM, Budinger GR, Perlman H. Flow cytometric analysis of macrophages and dendritic cell subsets in the mouse lung. Am J Respir Cell Mol Biol. Oct 2013;49(4):503–10. doi:10.1165/rcmb.2013-0086MA

52. Yu YR, O’Koren EG, Hotten DF, et al. A Protocol for the Comprehensive Flow Cytometric Analysis of Immune Cells in Normal and Inflamed Murine Non-Lymphoid Tissues. PLoS One. 2016;11(3):e0150606. doi:10.1371/journal.pone.01506

